# Constructing and Forgetting Temporal Context in the Human Cerebral Cortex

**DOI:** 10.1101/761593

**Authors:** Hsiang-Yun Sherry Chien, Christopher J. Honey

## Abstract

How does information from seconds earlier affect neocortical responses to new input? Here, we used empirical measurements and computational modeling to study the integration and forgetting of prior information. We found that when two groups of participants heard the same sentence in a narrative, preceded by different contexts, the neural responses of each group were initially different, but gradually fell into alignment. We observed a hierarchical gradient: sensory cortices aligned most quickly, followed by mid-level regions, while higher-order cortical regions aligned last. In some higher order regions, responses to the same sentence took more than 10 seconds to align. What kinds of computations can explain this hierarchical organization of contextual alignment? Passive linear integration models predict that regions which are slower to integrate new information should also be slower to forget old information. However, we found that higher order regions could rapidly forget prior context. The data were better captured by a model composed of hierarchical autoencoders in time (HAT). In HAT, cortical regions maintain a temporal context representation which is actively integrated with input at each moment, and this integration is gated by prediction error. These data and models suggest that sequences of information are combined throughout the cortical hierarchy using an active and gated integration process.

## Introduction

Events such as gestures, melodies, speech, and actions unfold over time, so we can only perceive and understand information in the present by integrating it with information from the past (Buonomano and Maass, 2009; Fuster, 1997; Kiebel et al., 2008). This process is complex because the world contains meaningful structure on scales ranging from milliseconds to minutes (Gibson et al., 1982; Poeppel, 2003; Zacks and Tversky, 2001): a series of phonemes makes up a word, a series of words forms a sentence, a series of sentences expresses an idea. How is the human brain organized to integrate information across multiple timescales in parallel?

We and others have argued that the human brain employs a distributed and hierarchical architecture for integrating information over time (Baldassano et al., 2017; Fuster, 1997; Hasson et al., 2015; Honey et al., 2012; Lerner et al., 2011; Runyan et al., 2017). The architecture is distributed because almost all regions of the human cerebral cortex exhibit temporal context dependence in their responses. The architecture is hierarchical because early sensory regions integrate over short timescales (milliseconds to seconds), while higher-order regions integrate information over longer timescales (seconds to minutes).

The timescale hierarchy is a highly reliable phenomenon with functional implications across the brain (Baldassano et al., 2017; Burt et al., 2018; Chaudhuri et al., 2015; Cocchi et al., 2016; Demirtaş et al., 2019; Watanabe et al., 2019), yet our models of the underlying information processing have remained phenomenological. What are the computations that integrate past and present information within the hierarchical networks of our brains? How are past and present information represented and combined within each stage of processing? What information is passed on to higher processing stages? We set out to answer these questions using a combined empirical and modeling approach.

To investigate how information is integrated over time, prior studies have measured the “processing timescales” of different brain regions. Processing timescales were quantified by comparing a brain region’s response to a stimulus at time *t* across various contexts, where the stimulus properties at time (*t-τ)* were altered. For example, Lerner et al. (2011) used functional magnetic resonance imaging (fMRI) to measure the neural responses to temporally manipulated versions of an auditory narrative (Figure 1A). They compared the neural responses during the original intact clip against the neural response during versions of the stimulus in which the ordering of words, sentences or paragraphs was scrambled. The authors observed that early sensory regions exhibited similar responses to the intact and scrambled audio; these early regions were said to have a short processing timescale, because their responses at each moment were largely independent of prior context. Moving toward higher-order cortices, such as temporoparietal junction, precuneus, and lateral prefrontal cortex, Lerner et al. (2011) observed different responses to the intact and scrambled input. In these higher-order regions, the response at one moment could depend on stimulus properties from 30 seconds or more earlier (Figure 1B). Overall, higher stages of cortical processing were said to have longer processing timescales, because their responses at time *t* were affected by properties of the stimulus from many seconds earlier (Figure 1C).

**Figure 1.**
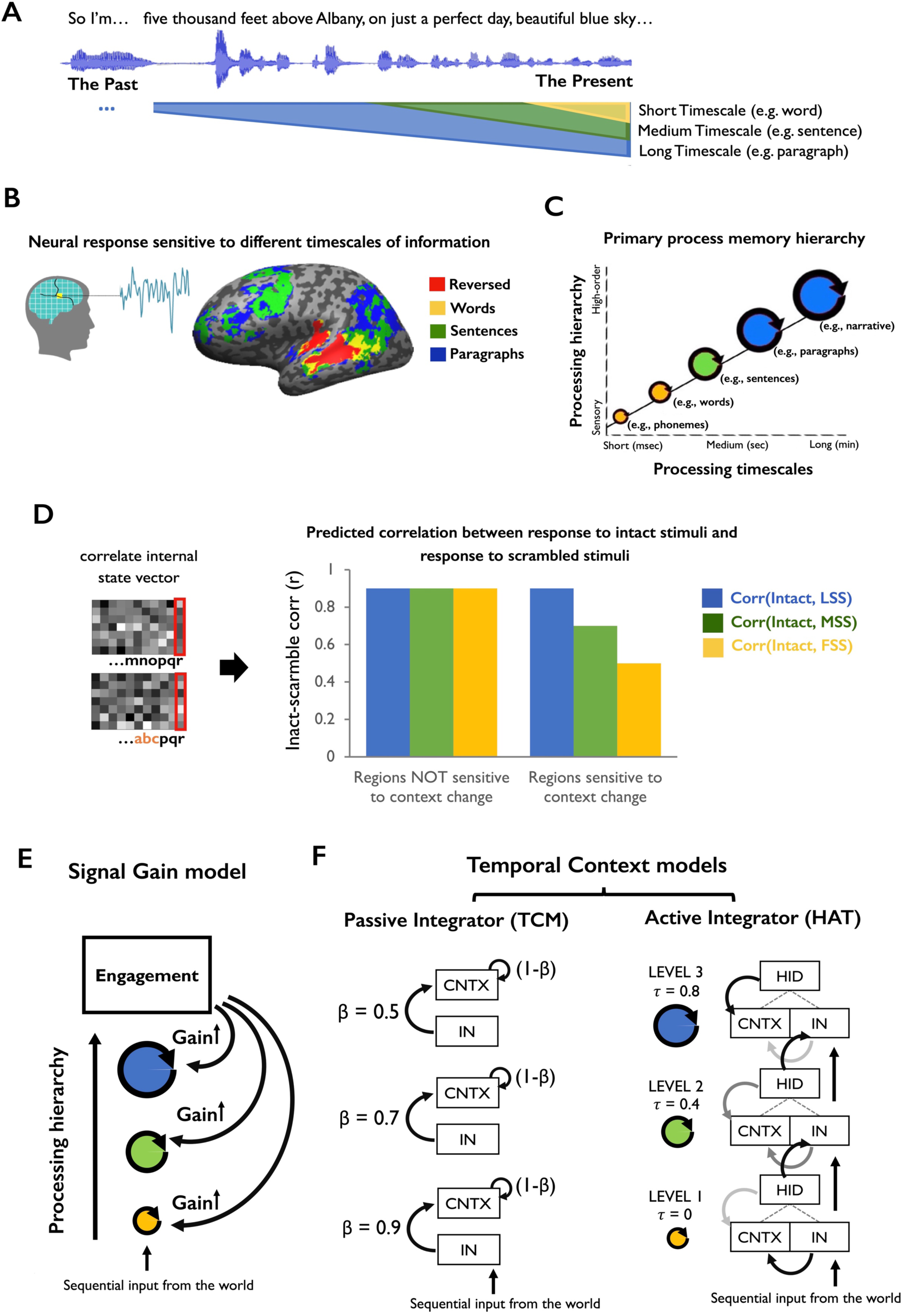
Computational models of distributed and hierarchical process memory. **(A)** Schematic of the experiment and results from Lerner et al. (2011). While in the fMRI scanner, participants listened to an auditory narrative, including an intact version and versions scrambled at different timescales (the scale of words, sentences and paragraphs). **(B)** The inter-subject correlation results indicated that lower-level regions (e.g. early auditory cortex) exhibited responses that were reliable across all stimuli, with little dependence on prior temporal context. By contrast, higher-level regions (e.g. TPJ and precuneus) exhibited responses that depended at each moment on long-timescale (tens of seconds) of prior context in the stimuli. **(C)** A schematic of the “process memory hierarchy”. Each circle represents a neural circuit which integrates information over different timescales. Lower-level regions (e.g. sensory regions) exhibit shorter integration timescales, integrating over entities such as phonemes and words. Higher-level regions (e.g. lateral and medial parietal regions) exhibited longer integration timescales, combining information on the scale of entire events (e.g. multiple paragraphs of text over 30-60 seconds). **(D)** Schematic of the predicted data when comparing the hidden representations of brain regions that are sensitive to temporal context on different scales. The dependent variable is the “intact-scramble correlation”, quantifying the similarity of neural response to the same input in different temporal contexts. **(E)** Schematic of a signal gain model for explaining the pattern of brain responses shown in panel D. The model posits that (i) overall response gain (or signal-to-noise ratio) increases for coherent stimuli, because they are more engaging, and that (ii) this effect is even larger in higher order regions. **(F)** (left) Schematic of a passive integrator model, TCM. The model posits that the new state of each brain region is a weighted sum of its old state and the new input. (right) Schematic of an active integrator model, HAT. Each region maintains a representation of temporal context. At each moment, representation of past and present information are combined to a simplified joint representation. LSS = long scale scramble, MSS = medium scale scramble, FSS = fine scale scramble. TCM = temporal context model. HAT = hierarchical autoencoders in time.

Our approach was to develop a set of computational models that could explain key features of existing data (i.e. Lerner et al., 2011) concerning the timescale hierarchy (Figures 1B, C), and then to empirically test predictions of these models by conducting new analyses on a new dataset. In the first section of this manuscript, we specify multiple models of hierarchical temporal integration, and test whether they can account for existing data. These computational models are important, because we provide a process-level specification of how information sequences are integrated within each stage of cortical processing and also what kind of information is sent between stages of processing – specifications that were lacking in prior descriptions of the timescale hierarchy. We also consider informal models that set out to explain the data using “stimulus engagement” rather than temporal context. Furthermore, we contrast temporal context models which employ passive integration mechanisms (temporal smoothing) and those with active (recurrent and gated) integration mechanisms.

In the second section of this manuscript, we report empirical data that can decide between the models. In order to test the predictions of each model, we apply a time-resolved pattern analysis to fMRI recordings of brain responses to naturalistic auditory stimuli. Specifically, we examine how each integration model can account for moment-by-moment changes of fMRI responses when two groups of participants hear the same natural auditory speech segments preceded by different contexts. We found that the fMRI responses gradually aligned over time across the two groups, when each group heard the same input preceded by a different context. The responses aligned earliest in sensory regions, but later and later in regions at consecutive stages of processing. The large-scale topography of these alignment patterns suggested that the time for building temporal context was hierarchically organized. This “hierarchy of context construction” provides direct evidence for representations of prior context across the cerebral cortex.

In the third and final section of this manuscript, we set out to test a prediction of the hierarchical autoencoders in time (HAT) model: that there are gating mechanisms for “forgetting” the temporal context in the cerebral cortex. Baldassano et al. (2017) observed rapid transitions in brain state (possibly reflecting event boundaries) while listeners perceived complex movies and stories. In those data, higher-level regions switched their activity pattern in alignment with a small number of long events, while lower-level sensory regions appeared to switch state at the transitions between many short events. The existing data do not determine whether the rapid switches reported by Baldassano et al., (2017) were (i) driven by rapid changes in the features of the input or (ii) reflecting a gating mechanisms for resetting temporal context at the start of a new event, which can reduce interference from irrelevant prior information (Reynolds et al., 2007). To directly test for context gating, we compared the “rate of integration” and “rate of forgetting” across all brain regions responsive to the real-life auditory narrative. Although higher order regions integrated information more gradually than sensory regions, we found that they did not “forget” prior information more gradually. The ability to integrate slowly and yet forget more rapidly provides cortical regions with flexibility in temporal integration: at appropriate moments, such as the start of a new event, regions can generate a response that depends less on prior context.

Altogether the models and data detail a framework for the computations implemented in the cortical hierarchy when humans are processing naturalistic sequences. The empirical findings of hierarchical context construction and context forgetting (i.e. temporal integration and separation) were consistent with the predictions of the HAT model. In this active integration model, individual cortical regions maintain a temporal context buffer, which can be updated to integrate past and present information, but which can also be overwritten upon arrival of surprising input.

## Results

We developed a set of models to explain prior measurements of neural responses to intact and temporally scrambled stimuli (e.g. Lerner et al., 2011, also Baldassano et al., 2017; Chen et al., 2017; Farbood et al., 2015; Hasson et al., 2008; Honey et al., 2012; Simony et al., 2016; Yeshurun et al., 2017). In particular, lower-level sensory regions should display context-invariant responses to the intact and scrambled narratives: the response to a particular input segment (e.g. a spoken sentence) in each of the scrambled conditions should resemble the response to that same segment within the intact condition. By contrast, higher-level cortical regions that are sensitive to temporal context change should show more divergent responses to a given input segment when the surrounding context is more scrambled, and thus we should observe a larger difference between the intact and scrambled conditions. The difference between intact and scrambled conditions is manifest as a reduced “intact-vs-scramble correlation”, where the correlation measures the similarity of responses to the same stimulus segment presented in the intact or scrambled context (Figure 1D). In sum, for a model to account for the hierarchy of context dependence it should capture two key phenomena:

(P1) lower processing stages of the model should be insensitive to context change (analogous to sensory cortical regions, Figure 1D, left bars);

(P2) progressively higher processing stages of the model should be increasingly sensitive to temporal context extending further into the past (analogous to the higher stages of cortical processing, Figure 1D, right bars).

We developed three computational models to account for these phenomena: a model based on engagement (signal gain model, Figure 1E), a model employing passive temporal integration (the temporal context model, TCM, Figure 1F, left), and a model employing active gated integration (HAT, Figure 1F, right).

### The signal gain model

It is possible to account for the empirical phenomena (P1 and P2, Figure 1D) without explicitly drawing on the notion of distributed temporal integration. Instead, one can offer an explanation based on “signal gain”, in combination with the qualitative notion of “engagement”. This model makes three reasonable assumptions: (i) when participants engage more deeply with a stimulus, the gain of their response to that stimulus increases relative to the noise level, and they produce a more reliable neural responses to that stimulus (Cohen et al., 2018; Dmochowski et al., 2012) (ii) participants are less “engaged” with scrambled stimuli than with intact stimuli, and (iii) the effects of engagement on neural reliability are larger in higher-order cortical regions. With these assumptions, the required pattern of results can be explained: first, sensory neocortex would be largely unaffected by engagement (and thus unaffected by scrambling prior context); second, higher order regions would respond less reliably to scrambled stimuli, and so their intact-vs-scramble correlations would also be decreased (Figure 1E, STAR Methods Section 1.2).

The signal gain model can explain data from scrambling experiment, without recourse to any neural representation of temporal context, and thus provides an important null model. We next considered models which do assume a representation of temporal context.

### The temporal context model (TCM)

We employ the temporal context model (TCM) to stand in for a general class of linear integrator models, in which past and present information are combined as a weighted linear sum. In TCM, the arrival of each new stimulus generates linear “drift” of an internal context variable, and this “drift” during sequence encoding can account for contiguity effects in later recall (Howard and Kahana, 2002; Howard et al., 2011). Can a linear combination of past and present information (passive integration) account for the empirical data from scrambling experiments?

#### Local processing in TCM

If we define the current context as CNTX(t) and the current input as IN(t), a simple form of the update equation for TCM is:

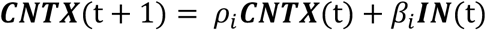

where *ρ_i_* and *β_i_*-are “drift” parameters that determine the proportion of new and old information in the updated context variable.

TCM employs a “passive” temporal integration because the drifting context, CNTX, is essentially an exponentially weighted running average of prior input. The external input is always linearly combined with the context, and so the relationship of new input and prior context (e.g. whether they are related or unrelated) does not affect the magnitude or form of the context update. Moreover, there is no mechanism for controlling (e.g. via gating) how much of the context is overwritten by new input at each moment – the proportional influence of the input depends only on the integration parameters, *ρ* and *β*, which are fixed, and could be adjusted by the status of learning (Sederberg et al., 2011). Thus, TCM employs a passive integration scheme in which the context variable is a weighted average of inputs.

#### Stages of processing in TCM

We created a hierarchical variant of TCM by testing multiple copies of TCM, each with different drift parameters (*ρ* and *β*). The drift parameters determine the proportion of influence of the current input. Thus, we increased *ρ* (and decreased *β*) to model higher stages of processing, thereby increasing the proportion of prior information that is retained at higher stages of the simulated hierarchy (Figure 1F, left; STAR Methods Section 1.1).

### The hierarchical autoencoders in time (HAT) model

Finally, we developed a model of neural sequence processing in which past and present information are actively integrated. To this end, we were inspired by TRACX2, a recurrent network model which can account for diverse human sequence learning behaviors (French et al., 2011). This is an active form of temporal integration, because new input and prior context are compressed into a simplified joint representation (Mareschal and French, 2017), and the output of the model from this joint representation depends on learning of what inputs typically arrive in which temporal contexts.

#### Local processing in HAT

We modeled individual processing stage as “autoencoder in time” (AT) units. At each time step, each AT unit attempts to generate a simplified, or compressed, joint representation (HID) of its current input (IN) and its prior context (CNTX, Figure S1, STAR Methods Section 1.3.1). The simplified representation (HID) is compressed, because IN and CNTX are jointly represented in a lower dimensional space. This simplified representation then becomes the context for the next input.

What happens when an unfamiliar or surprising input is presented to the HAT model? In this case, the network will be unable to generate a simplified joint representation of prior context and new input; the model overwrites its context variable (CNTX), and the current input (IN) instead becomes the context for the next time step. Intuitively, this should occur at boundaries between temporal chunks. For example, suppose we were to present the sequence of letters “o,c,e,a,n,v,i,e,w”, to the model, one by one. When the letter “n” is input, the model could form the joint representation “ocea+n” (ocean). However, when the letter “v” is input, the model cannot form a simplified joint representation for “ocean+v”. Because the letter “v” cannot be readily integrated with the prior context, the letter “v” becomes the initial context for a new subsequence (view). Thus, at the boundary between the two chunks, the context variable is “gated”, so that the letters in the new chunk (view) are separated from those in the previous chunk (ocean).

#### Hierarchical architecture in HAT

We constructed a hierarchical temporal autoencoder in time (HAT) model by stacking three levels of AT units (Figure 1F, right). The information flow between units is globally feedforward; there is recurrent signaling within each AT unit, but there is no feedback from a higher AT unit to lower stages. Higher order units possess longer intrinsic timescale *τ*, so their context is less readily influenced by the input from lower level units (Figure S2A, STAR Methods Section 1.3.2).

#### Information transmission in HAT

The information transmitted from a lower AT unit (stage *i*) to a higher AT unit (stage *i+1*) depends on the surprise *α* detected in the lower stage (Figure S2B, STAR Methods Section 1.3.2). When the lower level of the model is less surprised, it transmits more of its “compressed” representation HID_i_ as input IN_i+1_ for the next stage of processing. When the lower level is more surprised, it transmits more of its own input IN_i_ as input IN_i+1_ for the next stage of processing. This between-level gating mechanism helps the model to preserve the learned “chunks” from the lower levels to be processed by the higher levels.

### Testing computational models of hierarchical context dependence

We tested whether the signal gain model and the temporal context models (HAT and TCM) could capture the previously described phenomenon of hierarchical context dependence (Figure 1D). To measure context dependence in each model, we first trained the model by exposing it to sequences of input with temporal regularities. We then tested the internal representations generated by the models when the same (identical) inputs were presented within distinct larger sequences that provided different scales of coherent context. Further details of the modeling and more fine-grained model analysis are presented in STAR Methods Section 2.1-2.3 and Figure S3 & S4.

As expected, all three models – signal gain, passive integration (TCM) and active integration (HAT) – could account for the key empirical phenomena of hierarchical context dependence (P1 and P2, above). In all models, the lower levels exhibited high correlation between representations of intact and scrambled stimuli, regardless of the scale of scrambling. This demonstrates that these lower levels of the model are relatively insensitive to context change. In contrast, the higher levels of the models showed progressively more dependence on context: the correlation between representations of the intact stimuli and the long-scale scrambled stimuli remained high, but the representation of the intact stimuli no longer resembled the medium scale or fine-scale representations (Figure S3).

The HAT model did exhibit an important advantage over TCM, because its temporal integration was more selective for previously learned sequences (STAR Methods Section 2.3 and Figure S3). In addition, the preference of different levels for distinct timescales was more distinct, which better matches empirical data. We elaborate on these differences, as well as further tests of the HAT architecture, in Supplemental Text.

In summary, we developed three explanations for existing data concerning the cortical responses to intact and scrambled sequences of naturalistic input. We found that passive temporal integration (TCM) and active, gated integration (HAT) could all account for the basic empirical phenomena. However, to decide between the different forms of the temporal integration models, and to rule out the stimulus engagement accounts, it was necessary to collect more fine-grained measurements of hierarchical temporal integration.

### Measuring the Moment-by-Moment Construction of Temporal Context

We developed a time-resolved fMRI pattern analysis approach for measuring the responses to naturalistic auditory narratives with different shared context. This approach, applied here to a new fMRI dataset, provided three benefits. First, this analysis revealed a second-by-second picture of context-dependent processing in the human cortical hierarchy, going beyond previous maps based on the average response over minutes of processing. Second, the time-resolved data enabled us to test and rule out the “engagement” style explanations of the data. Third, the time-resolved data enable us to map both the timescales of information accumulation and the timescales of forgetting throughout the cortical hierarchy.

To understand the time-resolved analysis, consider a case in which two groups of subjects are exposed to the same ∼20 s segment of natural speech (e.g. sentence E), but this shared segment is preceded by different speech segments across the two groups (e.g. sentence C or sentence D, Figure 2A). In this setting we can ask: how similar are the neural responses within and across these groups, second by second, as they process the shared segment from start to end? At the very start of the sentence, the two groups share none of their prior context, but by the end of the sentence they share much greater amounts of prior context. Thus, context-based models (such as HAT and TCM) predict that the correlation *across* groups should start from zero (as the groups share no context at the start of segment) and ramp upward over time as more and more context become shared across the groups. At the same time, context-based models also predict that the correlation across participants *within* each of the two groups will, on average, be constant over time, because the participants in each group are hearing the same current input preceded by the same context. Thus, we can test the predictions of the context-based models by measuring the across-group and within-group similarity at each time step within each sentence.

**Figure 2.**
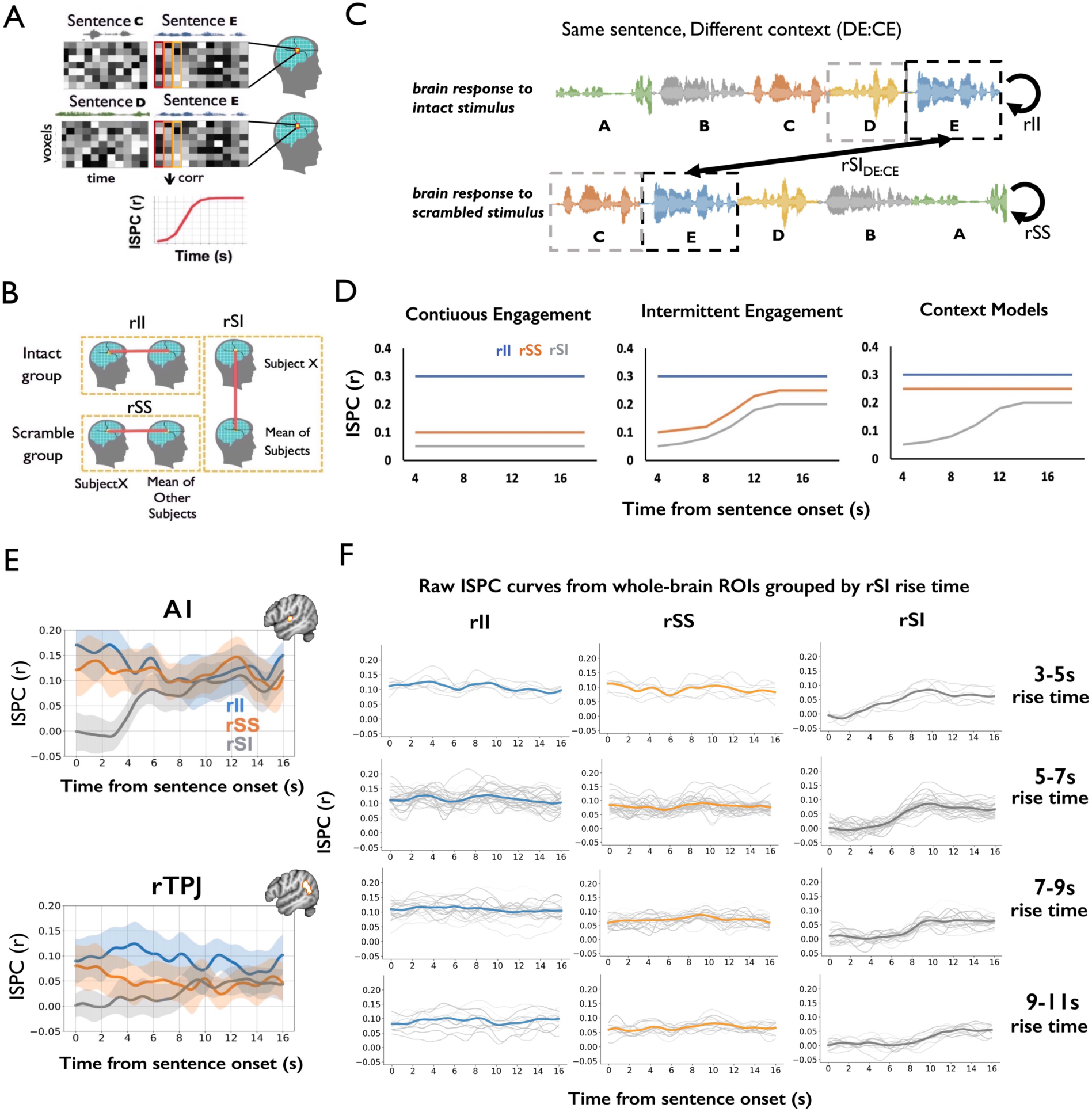
Gradual alignment of responses to a common stimulus preceded by different context. **(A)** For a specific sentence, inter-subject pattern correlation (ISPC) was measured by correlating the spatial pattern of activation at each time point across the two groups. **(B)** ISPC was calculated between one subject and the average of the rest of the subjects within the intact group (rII); or between one subject and the rest of the scrambled group (rSS); or across the intact and scrambled groups (rSI). **(C)** The ISPC analysis for the same sentences preceded by different contexts (DE:CE). In this example, sentence E followed sentence D for listeners in the Intact group, but it followed sentence C for listeners in the Scrambled group. **(D)** Predictions of ISPC generated from candidate models: two variants of the signal gain model, and two models that assume a representation of temporal context. **(E)** Average ISPC for all sentences in ROIs within an auditory (A1+) region and a right TPJ region. In A1+, the within-group rII and rSS curves showed similarly flat timecourses, while the cross-group rSI_DE:CE_ curve began at zero and then started to increase at ∼4 seconds. In the right TPJ, the within-group rII and rSS curves were again stable during the sentence processing. Crucially, the rSI_DE:CE_ curve was low at the beginning of the sentences but started to increase at around 8 seconds. Shaded area indicates a 95% confidence interval on individual rSI estimates. (F) Plots of the raw rII, rSS, and rSI_DE:CE_ curves for individual regions across the cerebral cortex, grouped according to their “rise time”. Curves for individual regions within that timescale grouping are in pale gray. The mean curve for each rise-time grouping is shown in thick blue (rII), orange (rSS), and gray (rSI_DE:CE_). The rII and rSS curves are stable over time, while the rSI curves consistently exhibit ramping. A1 = primary auditory cortex, rTPJ = right temporal-parietal junction, ISPC = inter-subject pattern correlation, rII = intact-intact ISPC, rSS = scramble-scramble ISPC, rSI = scramble-intact ISPC.

To empirically quantify the neural similarity within and across groups in a time-resolved manner, we calculate the inter-subject pattern correlation (ISPC, see STAR Methods Section 3). Three kinds of ISPC are calculated: similarity within the intact group (i.e. intact-intact correlation, rII), similarity within the scramble group (i.e. scramble-scramble correlation, rSS) and similarity across the intact and the scramble group (i.e. scramble-intact correlation, rSI) (Figure 2B, 2C).

Signal gain (or “engagement”) models and context models generate distinct predictions for how the set of similarity measures will evolve over time. A simple form of the engagement model (“continuous engagement model”) would propose that people are generically less engaged in the stimuli in the scramble condition. Thus, the rSS and rSI timecourses should be lower than the rII timecourse, and this effect should persist from the beginning to the end of each segment (Figure 2D, left). Alternatively, a more sophisticated form of this model could propose that there are time-varying changes in engagement. This “intermittent engagement model” proposes that people in the scramble condition are less engaged specifically at the beginning of each new segment (perhaps because they are confused by non-continuity at the boundary between unrelated segments). Thus, under this intermittent engagement model, both the rSS curve and rSI curve should be lower than the rII curve at the beginning of the segments, but then both the rSS and rSI curves should increase gradually as subjects in the scramble condition come to understand more of the segment they are hearing (Figure 2D, middle). The predictions of the engagement models differ from those of the context models (HAT or TCM). As noted earlier, the context-based models predict that rII and rSS should be stable over time, while the rSI curve should show a ramping phenomenon (Figure 2D, right).

In summary, the key prediction of the context models is that rSI should be low at the beginning the segment (due to the different prior contexts), and should rise later as more shared context is built across the two conditions, even while the rII and rSS curves are stable. Such an effect demonstrates that each group (intact and scrambled) is producing a reliable and stable response, but what is common across groups is changing, even while the input that the two groups receive is identical. Such an effect can only be explained by an effect of prior context on the neural response to the current input.

### Time-resolved Neural Similarity Data Match Predictions of Context Models

Overall, the time-resolved pattern-analysis results were consistent with the predictions of the distributed context models, and inconsistent with the predictions of engagement models. To start, we considered the curves of rII, rSS and rSI within one lower level region (an ROI near A1+, Figure 2E top) and one higher order cortical region (an ROI near the TPJ, Figure 2E bottom). In both of these example regions we observed that (i) the rII and rSS curves were essentially constant from the beginning to the end of a segment; and (ii) the rSI curve ramped upward over time, as the intact and scrambled groups were exposed to more and more common temporal sequence within the same segment. The ramping of the rSI timecourse is inconsistent with the continuous engagement model. The flatness of the rII curve is inconsistent with the intermittent engagement model. These data patterns (flat rII; flat rSS; ramping rSI) are preserved across the cerebral cortex (Figure 2F, Figure S7) when we broaden our analysis to a cortex wide atlas or ROIs (Schaefer et al., 2018). Thus, neither intermitted engagement models nor continuous engagement models can account for these data, even at a qualitative level.

We next examined how the temporal integration profile (rII, rSS and rSI) differed across regions. To illustrate the basic phenomenon, we examined the rII and rSS curves (within-group correlation) for one sensory region (A1+) and one higher-order region (right TPJ). In A1+, we found that rII and rSS were very similar to each other across the whole segments, suggesting that A1 showed highly reliable responses to the same segments in the two conditions in which the contexts are different (Figure 2E top). In rTPJ, on the other hand, the rII curve was significantly higher than the rSS (t(21)=2.83, p=0.007, t-test of mean rII and rSS values per segment, Figure 2E, bottom). This effect could reflect some influence of engagement; it can also be explained by the fact that the patterns in the intact stimulus are more familiar than the scrambled stimulus (more similar to prior experience).

The across-group correlation (rSI) ramped upward later in the higher order cortex (TPJ) than in sensory cortex (A1+). In A1+, the rSI timecourse begins to rise from zero ∼3 s after the segment- onset, while in TPJ the rSI timecourse rises from zero ∼8 s post-onset (Figure 2E). Importantly, the fact that the rSI = 0 at the onset of the segment does not necessarily reflect a neural context effect – instead, this reflects an effect of hemodynamics. In particular, the hemodynamics introduce temporal smoothing from the previous segment, carrying information over into the beginning of the current segment, even if the underlying neural response is unaffected by context. This HRF artifact makes it difficult to use BOLD imaging to estimate the shortest possible time at which temporal context effects operate. However, the hemodynamics cannot account for the ramping in TPJ beginning many seconds later than in A1+. Instead, the later rise time in TPJ points to a neural context effect, with a longer timescale in higher order regions. Having observed evidence of variable rise-times in these two specific regions, our next step was to map the timescales of context construction across the cerebral cortex.

### Moment-by-moment context analysis reveals a hierarchical organization

As predicted by hierarchical temporal context models (HAT and TCM), we observed a hierarchical organization of moment-by-moment context-dependent responses in the human cerebral cortex. The similarity of response across the intact and scramble groups (rSI) exhibited an increasing pattern in almost every ROI, but the latency of this ramping was greater in higher-order regions. Because the rSI measurement will not be meaningful when the response in the scrambled condition is unreliable, we restricted our analysis of the ramping to the 83 ROIs in which there was a reliable response to the scrambled stimulus (i.e. mean rSS > 0.06, see STAR Methods Section 3.7, Figure S5). We then used logistic fitting to quantify the timescale of ramping in each ROI. After measuring the rise time (i.e., the time that the curve reaches its half maximum), we excluded 4 ROIs that were not well-fit by a logistic function (Figure S6), resulting in 79 ROIs for further analysis. A direct visualization of the raw rSI timecourses in each ROI confirmed that the logistic fitting accurately captured the profile of the rSI curves (Figure S7). To further confirm that the latencies derived from the logistic fits accurately reflect the data, we grouped the ROIs based on the “rise time” (i.e. the time that the logistic fits reaching the half maximum, see Material and Methods Section 3): the 3-5s, 5-7s, 7-9s and 9-11s group.

These groupings revealed a consistent rSI ramping latency within groupings and distinct rSI ramping latency across groupings (Figure 2F). The second-by-second analysis revealed a “hierarchy of context construction” across the lateral and medical cortical surface, in which early auditory regions first arrive at a shared context-dependent response, followed by consecutive stages of the cortical hierarchy. Rise times of rSI curves became gradually slower from sensory cortex (with ∼ 4 s rise time) up toward higher order regions (rise times of 10 s or longer, Figure 3, top). The logistic curves extracted from five example ROIs along the auditory processing hierarchy were visualized, with ROIs drawn from A1, the superior temporal gyrus and the inferior parietal lobe (Figure 3, bottom). These rSI curves materially confirm the hierarchy of context construction within the auditory processing pathway: lower-level sensory regions (e.g. A1) quickly arrived at a shared response between intact and scrambled groups, while the inferior parietal regions (and regions in the medial parietal cortex) took longer to generate a response that is shared across the intact and scrambled groups.

**Figure 3.**
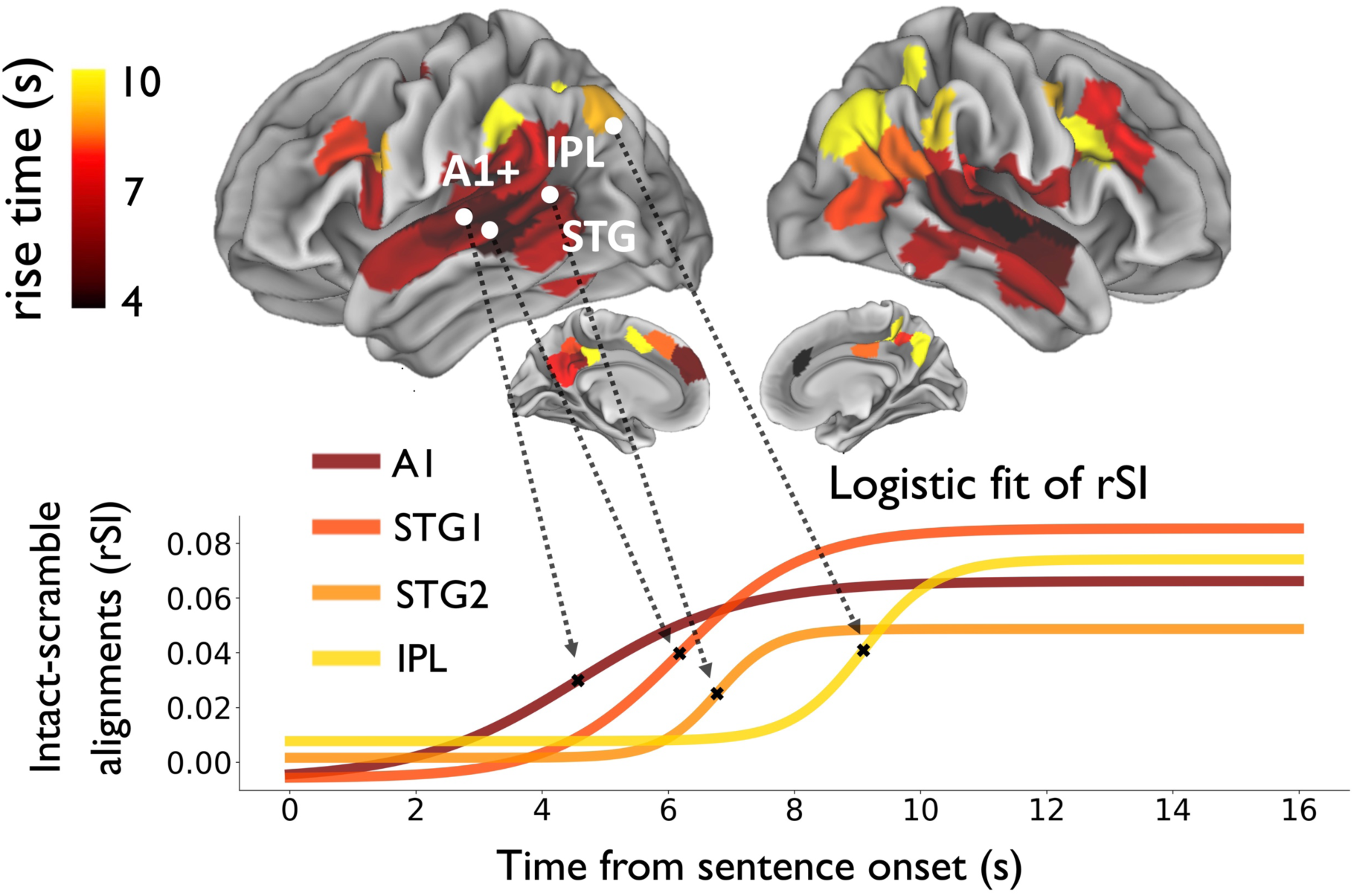
Hierarchical timescales of context construction across the human cerebral cortex. (top) Cortical map of the timescale at which neural responses align to a common input preceded by different contexts. Rise time is quantified as the time for each rSI_DE:CE_ curve to reach its half maximum value, based on logistic fitting. (bottom) Logistic curves are shown for four representative ROIs along the cortical hierarchy (from A1, to middle STG, to posterior STG, to IPL). A1 = primary auditory cortex, IPL = inferior parietal lobe, STG = superior temporal gyrus, rSI = scramble-intact inter-subject pattern correlation.

### Time-resolved analysis reveals an active context forgetting process

The time-resolved pattern analysis indicates that information is temporally integrated second-by-second throughout the cortex – but is integrating information over the past always desirable? For example, if the subject of a new sentence is unrelated to the verb of the previous sentence, then perhaps we might want to separate these pieces of information, rather than integrate them. Therefore, in addition to the mechanisms for integrating information over time, it seems desirable to be able to separate information from distinct events.

Establishing a new context will often involve forgetting a prior context, and so the processes of construction and forgetting may be connected (Figure 4A). Consider a setting in which two groups of subjects listen to the same auditory input preceded by different contexts. In the previous section, we showed that the two groups will gradually construct a shared mental context and will begin to respond in the same way to common input (Figure 4A, middle). But what happens when the common input ends? At this moment, the two groups begin to hear different inputs, but these different inputs are preceded by a shared context. We expect that the two groups should gradually “forget” the previously shared mental context, as they are exposed to distinct input, but the influence of prior context may persist for some time (Figure 4A, right).

**Figure 4.**
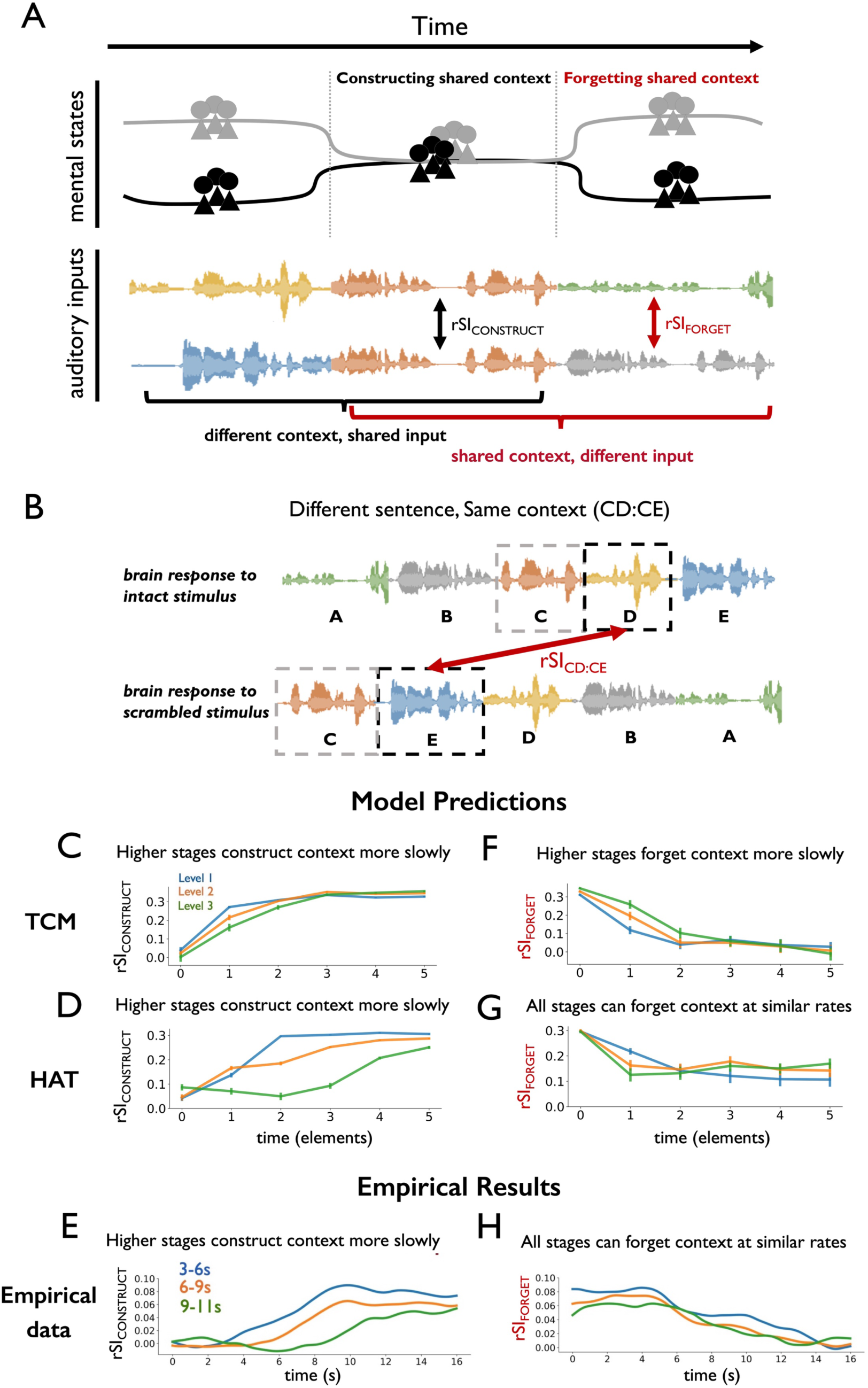
Distinct timescales of context construction and context forgetting. **(A)** Conceptual schematic of how mental states fall into and out of alignment as common and distinct input sequences are presented over time. Two groups of participants gradually construct a shared context when they listen to the same auditory input preceded by different contexts. As they construct the shared context their mental states (reflected in their neural responses) fall into alignment. Then, when the common input ends, they begin to process a distinct input preceded by a common context. As the distinct input is processed over time, participants forget their shared context. **(B)** Schematic of the inter-subject pattern correlation (ISPC) analysis for the different speech segments preceded by the same context (CD:CE). For example, in this diagram, segment D in the intact group and segment E in the scramble group were both preceded by segment C. **(C)** TCM simulation of rSI_DE:CE_ predicts a hierarchy of context construction. **(D)** HAT simulation of rSI_DE:CE_ predicts a hierarchy of context construction. **(E)** Empirical rSI_DE:CE_ results grouped by rise time, indicating a hierarchy of context construction, consistent with predictions of both TCM and HAT. **(F)** TCM simulation of rSI_CD:CE_ predicts that regions that construct context slowly will also forget context slowly, as temporal updating has a fixed time constant. **(G)** HAT simulation predicts that the timescale of context forgetting (rSI_CD:CE_) need not be slower in levels of the model that have longer timescales of context construction (rSI_DE:CE_). **(H)** Empirical rSI_CD:CE_ results grouped by rise time showed that regions in different levels of cortical hierarchy can forget context at a similar rate. This observation is inconsistent with the predictions of the TCM model, and more consistent with gated models such as HAT. TCM = temporal context model, HAT = hierarchical autoencoders in time, rSI = scramble-intact inter-subject pattern correlation.

How quickly can individual brain regions forget the previous shared context and start to build the new, different contexts? In the HAT model, it is possible to block the integration of prior information with new information using an *active forgetting mechanism*. Specifically, because of the surprise-driven gating in the HAT model (Figure 1E, Figure S2), the influence of prior context is reduced at moments of higher surprise. Conversely, in the TCM model, there is no active forgetting mechanism: temporal integration is passive, and the influence of past information on present responses depends only on the recency of the past information. More generally, higher-order regions are known to exhibit slower dynamics than sensory regions (Chaudhuri et al., 2015; Demirtaş et al., 2019; Honey et al., 2012; Murray et al., 2014; Ogawa and Komatsu, 2010; Stephens et al., 2013) and the presence of intrinsically slower dynamics provides a passive form of integration. It is therefore critical to test for evidence of a mechanism for individual regions to actively and flexibly adapt the integration timescale.

For the construction of temporal context, both the hierarchical TCM and HAT model predict the basic empirical results (Figure 4E) in which higher-level regions construct new context (ramp up in rSI_DE:CE_ or rSI_CONSTRUCT_) more slowly than sensory regions (Figure 4C & 4D). Indeed, such predictions can even be generated by variants of the HAT model with modified or eliminated “context gating” mechanisms (see Figure S8A-S8C). However, in models such as TCM, in which information is integrated with a fixed time constant, the rate of accumulating new information must be directly proportional to the rate of forgetting old information. This leads to a testable prediction: if temporal integration within each brain region only relies on a fixed time constant, then regions which integrate information more slowly (i.e. higher order regions) should also forget prior information more slowly.

To test this prediction, we designed an analysis which examines how rapidly each brain region “forgets” shared context. We quantified the “rate of forgetting” by measuring the similarity of neural responses as participants were processing different input preceded by a shared prior context (i.e., rSI_FORGET_ or rSI_CD:CE_, Figure 4B). The TCM model predicts that regions which slowly integrate information would also slowly forget prior context (Figure 4F). In contrast, the HAT model predicts that although higher-level regions integrate information over longer timescales than lower-level regions, they do not necessarily forget prior context at a slower rate, because they have access to an active gating mechanism (Figure 4G). The three HAT variants with limited context gating mechanism generated predictions that were more similar to the TCM model: higher-level regions forgot context more slowly than lower-level regions (Supplemental Text, Figure S8D-S8F).

Finally, to test the different predictions regarding the rate of context forgetting, we grouped brain regions according to their rise time in accumulating new information (rSI_CONSTRUCT_ or rSI_DE:CE_) and then visualized the rate at which they forget prior information (rSI_FORGET_ or rSI_CD:CE_). Crucially, we observed that the rSI_FORGET_ curves decreased at a similar rate, regardless of whether the corresponding rSI_CONSTRUCT_ curve had a fast or a slow rise time (Figure 4H). Furthermore, we observed no reliable correlation between rise time and fall time in individual ROIs (r = −0.1, p=0.5, Figure S9, fall time defined analogously to rise time of the logistic curve).

The analysis of forgetting rates results rules out models which employ a fixed time constant for temporal integration. Instead, the results are more consistent with models such as HAT, which possess an active mechanism for overwriting prior context at the transition to an unexpected input. Gating has been employed in single-module memory context models (e.g. CMR, Polyn et al., 2009) and in machine learning theory (e.g. LSTMs, Hochreiter and Schmidhuber, 1997), but these results suggest that gating may be employed in a distributed and hierarchical manner in the brain.

An active gating mechanism, acting in a distributed manner across the cortex, could provide an explanation for the observation of rapid shifts in brain state (“event segmentation”) occurring in a hierarchical fashion across the cortex (Baldassano et al., 2017). More generally, the data indicate that the temporal integration of information in the neocortex is an active and gated process, which cannot be explained solely by variation of a fixed time constant across regions (Chaudhuri et al., 2015; Honey et al., 2012; Murray et al., 2014; Stephens et al., 2013).

## Discussion

This study investigated how the human cerebral cortex flexibly integrates information over time. We implemented models of both passive and active temporal integration processes, and showed how they could account for existing data on temporal processing. These models predicted a “hierarchical context construction” phenomenon, which we then tested using a time-resolved inter-subject pattern correlation (ISPC) analysis. We observed the predicted phenomenon across the later cerebral cortex of the human brain: when two listeners heard the same sentence preceded by different contexts, their neural responses were initially different, but gradually fell into alignment. Sensory cortices aligned most quickly, followed by mid-level regions. Higher-order cortical regions aligned last, sometimes requiring more than 10 seconds of common input before the responses aligned across contexts. Our ISPC approach also enabled us to measure the rate at which prior context was “forgotten”. These forgetting analyses showed that, despite their long integration timescales, higher order cortical regions could sometimes rapidly forget prior context. These data are incompatible with passive integration models in which past and present information are linearly mixed. Instead, the data point to an active integration process in which past and present information are compressed into a simplified joint representation. Moreover, the influence of past information appears to be gated, as it does not have a fixed timescale, and our modeling suggests that such gating can be controlled by a local prediction error.

The theory of hierarchical timescales in the cerebral cortex is increasingly influential across cognitive, systems and clinical neuroscience (Baldassano et al., 2017; Burt et al., 2018; Chaudhuri et al., 2015; Cocchi et al., 2016; Demirtaş et al., 2019; Fuster, 1997; Hasson et al., 2015; Kiebel et al., 2008; Murray et al., 2014; Runyan et al., 2017; Scott et al., 2017; Watanabe et al., 2019). Studies have measured both “integration timescales” and “dynamical timescales” of brain regions. The “integration timescale” measures the timescale on which prior information affects the response to new input. By contrast, the “dynamical timescale” is a measure of how quickly or slowly population dynamics vary in a given brain region. Both ECoG data (Honey et al., 2012) and fMRI data (Stephens et al., 2013) have demonstrated a direct correlation between integration timescales and dynamical timescales. Moreover, single unit data in macaques (Murray et al., 2014; Ogawa and Komatsu, 2010) and optical imaging in mice (Runyan et al., 2017) revealed that dynamical timescales are longer in higher order areas. Thus, consecutive stages of cortical processing are expected to exhibit both (i) slower population dynamics and (ii) a longer window for integrating past information with new input.

There have been advances in mapping and modeling fast and slow cortical dynamics, but it remains uncertain how these dynamics relate to the functional and computational principles of hierarchical temporal integration remain. Accounting for differences in dynamical timescales improves models of large-scale brain dynamics at the individual subject level (Chaudhuri et al., 2015; Demirtaş et al., 2019). Strikingly, the gradients of timescales in brain dynamics correlate with gradients of myelin density (Glasser and Van Essen, 2011), gene transcription (Burt et al., 2018) and anatomical connectivity (Margulies et al., 2016). Moreover, dynamical timescales are altered in autism, and are correlated with core symptoms of the disorder (Watanabe et al., 2019). Thus, variations in timescales are a major feature of large-scale brain dynamics, but it remains unclear how they relate to the functional processing of information arriving from the world. This motivated us to ask: what computations are employed to combine past and present information at each stage of cortical processing? And how does this computational process relate to the gradient of dynamical timescales?

We developed both passive and active models of temporal integration, and tested whether they could account for the empirical phenomena described above. Both the TCM model (which passively and linearly mixes past and present) and the HAT model (which actively compresses and gates past and present information) could successfully account for the key phenomenon of hierarchical temporal integration (Lerner et al., 2011): in both models, the representations at higher stages were more affected by a longer window of input history. However, it was also possible to account for aspects of the empirical data via variations in stimulus engagement, rather than timescales of processing (Figure 1E). Therefore, we used the TCM and HAT models to predict a more precise and time-resolved data pattern (“hierarchical context construction”) that could not be explained by engagement effects.

We empirically verified the prediction of hierarchical context construction, using a new method for parametrically mapping context effects in cortical responses on the scale of seconds. Prior studies inferred processing timescales by comparing the responses of a brain region across different temporal permutation of a stimulus: e.g. comparing the responses across word-scrambled or sentence-scrambled stimuli, one might conclude that a region was integrating information at the scale of words or sentences. Here, by combining inter-subject pattern correlation approach and curve fitting, we measured the second-by-second influences of new input and prior context on the neural responses produced to natural spoken sentence. This analysis confirmed the predicted phenomenon of hierarchical context construction (rSI curves, Figure 2F, Figure 3, Figure S7): when two participants heard the same sentence preceded by different contexts, their early sensory cortices aligned earliest, followed by secondary cortices, while some higher order regions did not align until participants shared 10 seconds of continuous common input.

Hierarchical context construction (Figure 3) cannot be explained by differences in stimulus engagement or attention. Because scrambled stimuli are expected to be less engaging than intact stimuli, there was some ambiguity in the interpretation of prior evidence for distributed and hierarchical temporal integration (e.g. Lerner et al., 2011). In particular, higher order regions might provide an unreliable response to scrambled stimuli simply because the scrambled stimulus is less engaging. This engagement effect could lead to differences in representation of the intact and scrambled stimuli, which might be taken as evidence for a distributed context representation, when in fact the scrambled stimulus was not being represented reliably at all. The time-resolved ISPC approach (Figure 2B, C) showed that the correlation across groups (rSI, intact vs. scrambled) ramped upward as the two groups were exposed to the same stimulus (Figure 2F, rSI), and this occurred even while the correlation within each group (rII, rSS) did not change (Figure 2F, rII and rSS). In other words, higher order regions produced reliable responses to both the intact and scrambled stimuli, stably from the start to the end of a sentence. For that same sentence, as the two groups heard more common input, second-by-second, their responses to the identical input gradually became more similar. This pattern of results cannot be explained by engagement effects. Instead, these patterns suggest the existence of a distributed and multi-scale representation of prior context, which affects the neural response to input at the moment.

Although different cortical regions build temporal context at different rates, corresponding to the cortical hierarchy, those regions do not “forget” the context at the same rate as they build the context (Figure 4E, H). This implies the existence of a mechanism for flexibly altering how the past influences present responses. Linear integrator models (such as TCM) lack such flexibility: the rate of constructing context and the rate of forgetting context are both inversely related to a fixed parameter, *ρ*, and so the past and present information are linearly mixed in the same way regardless of their content. By contrast, gated context models such as HAT can account for different rates of forgetting and constructing context. This is primarily due to how the local context is “gated” by prediction error. In HAT, if prior context can be successfully compressed with new input, then the context is preserved, but if prior context and new input are incompatible (prediction error), then the context is overwritten. This surprise-driven gating mechanism is consistent with evidence for pattern violations being signaled independently at multiple levels of cortical processing (Bekinschtein et al., 2009; Himberger et al., 2018; Wacongne et al., 2011).

A context gating mechanism is important for clearing out irrelevant context at the boundaries between chunks or events (DuBrow et al., 2017; Reynolds et al., 2007). Sequences of information cannot be integrated indiscriminately: the subject of a new sentence is not necessarily related to the verb of the previous sentence. Indeed, recent data suggest that almost all stages of cortical processing are sensitive to event structure, with sensory regions changing rapidly at the boundaries between shorter events (e.g. eating a piece of food) and higher order regions changing at the boundaries between longer events (e.g. having an entire meal) (Baldassano et al., 2017). The data from Baldassano et al. (2017) revealed that cortical states could change rapidly at multiple levels of processing, but they did not determine whether this effect was due to gating of a context representation. Because the immediate stimulus and its preceding context always covaried, it was uncertain whether rapid cortical state changes reflected rapid changes in input, rapid changes in context representations, or both. Here, by separately controlling current input and prior context, we demonstrated that with the sharp event boundaries introduced in our stimuli, the local context could be actively overwritten at those boundaries. Additionally, we showed that such context gating can occur even at the highest stages of cortical processing. Our data suggest that when neural state shifts rapidly at event boundaries, this effect is not simply due to changes in the current input, but is due to a change in the influence of a context representation. The gating of context may operate via an immediate prediction error, as in the HAT model, or via a more diffuse breakdown of temporal associations (Schapiro et al., 2013)

At a computational level, gating of context is useful not only for representing information, but also for learning sequential structure. Gated neural networks have been widely applied in sequence learning models to capture long-range temporal dependencies (Hochreiter and Schmidhuber, 1997). Combining both the gated neural network approach and probabilistic inference to model human ability in event segmentation and generalization, the structured event memory (SEM) model successfully produces human-like event segmentation and identifies event schemata in naturalistic video data (Franklin et al., 2019). Moreover, gating is a broadly useful process in biological models of working memory, not only for maintenance of information but also for flexible updating and integration (Heeger and Mackey, 2018). The gating in the HAT model is less flexible than in machine learning architectures (e.g. long-short-term-memory networks, LSTM). This is because while gates in neural networks (such as forget gates in LSTMs or update gates in GRUs) can be triggered by arbitrary states anywhere else in the network, the gating in HAT is determined entirely by a local prediction error. However, the gating in the HAT model is sufficient to enable higher stages of processing to maintain stable context over long periods, while also being able to quickly update at the boundary of a new “chunk”.

What is the functional difference between “passive” versus “active” temporal integration? Here we define passive integration as occurring when the combination of new and old information is a linear mixture (or weighted sum) of past and present activity. For example, in perceptual decision making, a vector of new evidence is thought to be linearly combined with prior evidence at each time step (Mazurek et al., 2003; Townsend and Ashby, 1983). By contrast, an active integration mechanism is one in which the integrated representation (A and B) depends on priors concerning the co-occurrence of A and B (Carpenter and Grossberg, 1987). For example, when we hear the sounds “basket” and “ball”, the ability to generate the chunk “basketball” depends on our knowledge and experience with language. In the HAT model, learning to generate such chunks occurs via the training of a temporal autoencoder. Thus, the learning of sequential structure in HAT is a process of learning to combine prior context and new input into a compressed (lower dimensional) representation (Mareschal and French, 2017). The compression of the past and present is only possible if sequential regularities can be identified (e.g. the regular sequence of “ball” following “basket”). Thus, the HAT model had difficulty integrating random, unfamiliar sequences of information because it did not contain the appropriate knowledge to compress past and present, and was continually surprised by novel combinations (Figure S4). This content-specific impairment in HAT may be analogous to the difficulty a person experiences when trying to integrate sequences of words in an unfamiliar language.

Our computational approach was inspired by the multilevel neurocognitive models of Botvinick, (2007) and Kiebel et al. (2008). In both of these models, higher stages of cortical processing learned or controlled temporal structure at longer timescales. More generally, in machine learning, multi-scale architectures have been proposed for reducing the complexity of the learning problem at each scale, and for usefully representing a multi-scale environment (Chung et al., 2016; Jaderberg et al., 2019; Mozer, 1992; Schmidhuber, 1992). In neuroscience, multiple timescale representations have been proposed for learning value functions (Sutton, 1995), for tracking reward (Bernacchia et al., 2011), and for perceiving and controlling action (Botvinick, 2007; Paine and Tani, 2005). Moreover, the concept of temporal “grain” is influential in theories of hippocampal organization (Brunec et al., 2018; Momennejad and Howard, 2018; Poppenk et al., 2013; Shankar et al., 2016) and cortical organization (Baldassano et al., 2017; Fuster, 1997; Hasson et al., 2015; Lü et al., 1992; Wacongne et al., 2011). Consistent with hierarchical timescale models, we find that more temporally extended representations are learned in higher stages of processing, where dynamics change more slowly. We provide additional empirical constraint on future models by revealing the moment-by-moment time-course of context construction across the lateral cerebral cortex (Figure 3, 4E), and by demonstrating that even slowly-evolving context representations at higher levels can be rapidly updated at event boundaries (Figure 4H).

Our results are consistent with the finding that recurrent neural networks can provide a better prediction of neural responses than feedforward models (Shi et al., 2018), especially for the later component of the neural response beyond the feedforward sweep (Kar et al., 2019). Most relevant to the present work, Shi et al., (2018) showed that adding recurrence to a convolutional neural network improved its performance as an encoding model for visual pathway responses to a movie stimulus. Only neural responses to the intact stimulus were modeled in that study: however, higher stages of the model learned more context-dependent representations, and these produced a better model of neural response at higher stages of visual processing, providing a computational account of process memory (Hasson et al., 2015). More generally, many recurrent neural networks may be able to instantiate the empirical properties of hierarchical context construction (Figure 3, 4E) as well as gating (forgetting) of context (Figure 4H). In ongoing work, we are comparing neurobiologically plausible architectures that are consistent with the construction and forgetting phenomena we have described.

Methodologically, the pattern-correlation method used here provides several practical advantages for measuring integration timescales. First, we showed that it can be used to measure timescales of context forgetting in addition to context construction (Figure 4B). Second, the method is efficient: if reference data exists for the responses to the intact stimulus, then an rSI curve can be computed in a single participant using one presentation of one scrambled stimulus. Third, the rSI curve provides a profile of how context influence varies over time; we focused on rise-times in this study, but the asymptote and slope of the rSI curve can also constrain quantitative models of temporal integration.

## Limitation and Future Directions

For reasons of parsimony, we chose to model temporal integration using only within-layer recurrent connections (i.e. without inter-regional feedback). Of course, there is rich anatomical and functional reciprocity in the brain (Bastos et al., 2012; Markov et al., 2013; Sporns et al., 2007) and many models of cortical function emphasize the important of feedback and prediction (Friston and Kiebel, 2009; Heeger, 2017; Heeger and Mackey, 2018; Rao and Ballard, 1999). It is not clear which expectation effects in sequential processing rely on top-down predictions from high-level representations, as opposed to more local recurrent integration or facilitation mechanisms (e.g. Ferreira and Chantavarin, 2018). Although long-range feedback connections are undeniably essential for some brain functions (e.g. attentional control and imagery), and models with (weak) long-range feedback connections could also account for our data, the local recurrence of the HAT model was sufficient to account for the temporal integration processes we measured here during narrative comprehension.

In future of extensions of this work, we will train HAT variants on linguistic corpora, and use these to generate context-aware encoding models of the neural response to complex language (e.g. Jain and Huth, 2018). Two other important questions for future work will be (i) whether gating of past context is a binary or graded, depending on the magnitude of local prediction error; and (ii) whether context gating can occur entirely independently across distinct levels of processing.

To recap, the results of this manuscript derive from a combined modeling and empirical approach. First, we developed computational models of hierarchical temporal integration. We then tested the predictions of these models using new methods for mapping the timescales of integration and forgetting across the human cerebral cortex. We showed that brain regions align, second-by-second, in a hierarchical gradient, when they are exposed to a common input preceded by distinct contexts. We ruled out alternative explanations based on engagement, and empirically established that cortical regions maintain a distributed representation of prior context. Finally, by analyzing the forgetting timescales of cortical regions, we ruled out temporal integration models with fixed time constants. Our models and data provide concrete constraints for models of brain function in which memory is inherent across perceptual and cognitive function (Buonomano and Maass, 2009; Frost et al., 2015; Fuster, 1997; Hasson et al., 2015; McClelland and Rumelhart, 1985; Shi et al., 2018), and our computational modeling points to general principles – active integration and gating – that are used in temporal information processing across the cortical hierarchy.

## Supplemental Information

Supplemental Information includes nine figures and supplemental text.

## Acknowledgment

The authors gratefully acknowledge the support of the National Institutes of Mental Health (R01MH119099 to CJH; R01 MH111439-01 subaward to CJH); the NVIDIA Corporation (GPU Grant); the Sloan Foundation (Research Fellowship to CJH); and the Government of Taiwan (Graduate Scholarship to HSC). We thank Janice Chen, Ken Norman, Anna Schapiro, Hongmi Lee, Uri Hasson and Jinhan Zhang for their insightful feedback and discussion.

## Author contributions

(CRediT taxonomy): Conceptualization: HSC and CJH; Methodology: HSC and CJH; Formal Analysis: HSC and CJH; Investigation: HSC and CJH; Resources: HSC and CJH; Writing – Original Draft: HSC and CJH; Visualization: HSC; Supervision: CJH.; Funding Acquisition, HSC and CJH.

## STAR Methods

### 1. Computational Models of Hierarchical Temporal Integration

#### 1.1 The Temporal Context Model (TCM)

TCM can successfully account for human sequence encoding and retrieval behavior, using the concept of a drifting internal context (Howard and Kahana, 2002). TCM employs a feature buffer which contains a vector of features of item processed in the sequence stream, and it employs a context buffer (*CNTX*) composed of a “temporal context vector”. A matrix M^FT^ maps stimulus features to their corresponding representation in the context space, thus generating an “input vector” *IN* which is used to update the internal context. The temporal context updated by adding the mapped input ***IN***(*t*) to the prior context ***CNTX***(t) (equation 1).

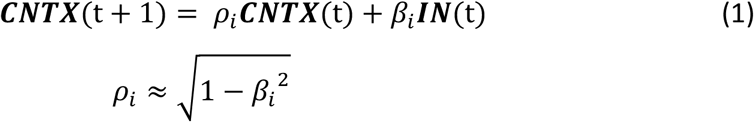

Note that we are focusing here on how TCM functions during encoding, and do not consider the “retrieved context” mechanisms that are fundamental to the model’s original purpose. In this sense, our main concern is with linear integrator models (e.g. Estes, 1955; Mensink and Raaijmakers, 1989; Murdock, 1997), and TCM is presented as a well-known example of this class. There are semantic variants (Polyn et al., 2009) and predictive variants (Howard et al., 2011) of TCM which should have a greater capacity to distinguish natural and unnatural sequence continuations during encoding.

#### 1.2 The Signal Gain Model

We implemented a simple signal gain model by using the same training and testing procedure as the temporal context model (See Section 2.1), while eliminating any effect of temporal context on the internal representation. In particular, we set *ρ* = 0 in Equation (1). To simulate scrambling effects across regions, we decreased the signal-to-noise ratio in the model for higher processing stages or finer scrambling conditions. In particular, the noise amplitude, *σ*, was re-scaled in the following simple model:

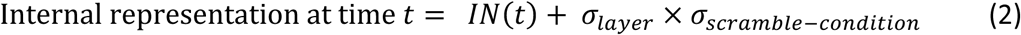

where we set *σ*_DEFGH_ = 0 in Layer 1, *σ*_DEFGH_ = 0.05 in Layer 2, and *σ*_DEFGH_ = 0 in Layer 3; and where we set *σ*_scramble-condition_ = 0.1, 0.5 or 0.9 for the paragraph-level, sentence-level, and word-level scrambling conditions.

#### 1.3 The HAT Model

##### 1.3.1 Local processing unit: the autoencoder in time (AT) module

Each local processing stage in HAT is an autoencoder in time (AT) module. This AT module was adapted from the influential TRACX2 model for modeling human statistical learning and sequence learning behavior (Mareschal and French, 2017). Each AT module has three layers. There is an input layer (consisting of a concatenated input unit, IN, and a context unit, CNTX); there is a hidden layer (HID) which stores the compressed representation of the input layer; and there is an output layer storing the reconstruction of the input layer from the compressed HID representation (Figure S1). During training, the model will learn good internal (i.e. HID) representations of the [CNTX, IN] pairings that frequently co-occur. At the end of training, it should be able to accurately reconstruct “chunks” of input-and-context from a compressed internal (HID) representation.

In the AT module, information from the world is presented as a stream of symbols, one symbol at a time. For every time step of the model, the current input symbol, *S_t_*, from the sensory environment is represented as a 1-by-N one-hot vector, where one scalar value is 1 and all others are −1. This new input vector is mapped to the IN bank at each time step. The prior context stored in the model is represented as another 1-by-N vector, which is stored in the CNTX bank.

A single time-step of the model proceeds as follows (please refer to Figure S1):

***A.*** *The model is initialized with two consecutive stimuli (S_t-1_, S_t_) in the CNTX and IN banks (CNTX_t_ = S_t-1_, IN_t_ = S_t_).*

***B.*** *Activity is propagated forward from the input and context banks (jointly of length 1-by-2N) to the hidden bank (1-by-N) via an affine transformation followed by a hyperbolic nonlinearity.*

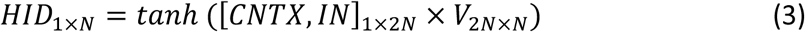

*This results in the activation of the hidden units in the model, and, following another linear-nonlinear transformation, the output nodes. Specifically, the auto-associative component of the model is implemented via the compressive transformation from the input and context* (*1-by-2N) to the hidden units, HID_t_ (1-by-N). A weight matrix V (of size 2N x N) contains the synaptic weights that transform the input layer to the hidden layer in the compression stage. A second weight matrix, W (of size N x 2N), is then right-multiplied with the hidden layer vector, HID_t_, to approximately reconstruct the CNTX and IN banks of output units:*

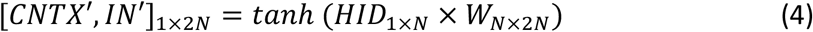

***C.****The objective is to make the reconstructed [CNTX’, IN’] as similar as possible to the input [CNTX, IN]. Therefore, an auto-associative error Δ is generated as the absolute difference of the input and output layer (the difference of the veridical and reconstructed representations):*

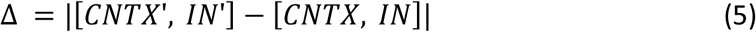

***D.*** *The “surprise” parameter, α, is calculated as the maximum value of Δ, multiplicatively scaled by a parameter k:*

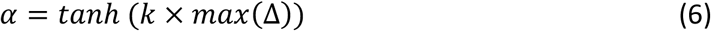

*α is taken to indicate the “surprise” or “familiarity” that the model experiences in response to the combination of the current context, CNTX, and the current input, IN. When k is larger, the average amount of surprise (magnitude of α) is increased, and IN makes a larger contribution to the CNTX variable at the next time step*.

*The CNTX bank is updated as a linear mixture of IN(t) and HID(t), weighted by the surprise parameter, α:*

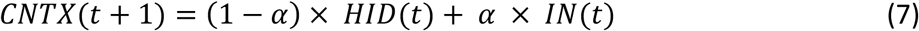

*If α is large, the model has not learned a good HID representation for accurately reconstructing IN_t_ and CNTX_t_, and so the CNTX_t+1_ bank will be overwritten input IN_t._ If α is small, the model has learned a good compressed joint representation, HID_t_, and this compressed representation becomes the context for associating with the next sequential input IN_t+1_*.

*This completes one update of the model. The process continues from **A**, and IN_t+1_ is set equal to the next element in the sequence, S_t+1_*.

Within each local processing unit, the transformation matrices (matrix V mapping from input layer to hidden layer, and matrix W mapping from hidden to output layer, Figure S1) are then adjusted via backpropagation to minimize the norm of the auto-associative error vector Δ. The backpropagation weight updates are performed incrementally, one training exemplar at a time. Backpropagation is entirely local to each processing unit (it is not performed end-to-end across the entire network, even when AT units are stacked).

As the auto-associative error gradually decreases, the model learns to capture the sequential structure of the input stream, by learning good internal representations of [CNTX, IN] sequences. The model can also detect the event boundaries occuring in the sequence. At event boundaries, the model will be unable to generate an accurate compressed representation of [CNTX, IN], and will generate a large error *α*. This large error will then bias the model to overwrite its prior context (from the old event) with its current input (from the new event).

In summary, the AT module exhibits three important features: (1) prior context is preserved in the CNTX bank; (2) the updating / overwriting of prior context is gated by an auto-associative error Δ which is summarized in the “surprise” parameter, *α*; and (3) the model minimizes its auto-associative error Δ by learning the statistical relationships between prior context and new input. We hypothesize that each stage of processing in the cortical hierarchy exhibits these three functional properties. The HAT model is thus composed of a stack of AT modules, each with these functional properties.

##### 1.3.2 Stacking the AT modules: Hierarchy of Autoencoders in Time

We employed a HAT model with three levels (Figure 1E). Each level is an AT module. The information flow in HAT is globally feedforward with local feedback: there is no backward information flow from AT module *i+1* to AT module *i*, but each AT module has recurrent feedback from its own past state.

Information processing in the HAT architecture possess two key features:

###### Local context is updated by level-specific constant *τ* and local surprise *α*

Figure 2B illustrates the structure of each AT module in HAT. As described in section 1.1, the AT module transforms the input and context [CNTX, IN] into a compressed internal representation, HID, and the model then attempts to reconstruct the [CNTX, IN] pairing from this lower-dimensional internal representation. The local context in each level unit is updated by a combination of HID and IN, modulated by a level-specific time constant *τ* and local surprise *α*, respectively (Figure S2A). If *τ* is larger than *α*, the model tends to preserve more context from HID; if *α* is larger than *τ*, the model tends to overwrite the context using the current input IN, as the equation illustrates:

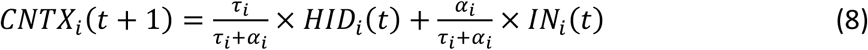

Note that in the full HAT model implementation reported in our simulations, we employed Equation (8) rather than the simpler Equation (7) which describes a single AT module.

To capture the assumption that higher-level regions process information over longer timescales while lower-level regions process information over shorter timescales, we set *τ* equal to 0.8 for the top level, 0.4 for the middle level and 0 for the bottom level of the 3-level HAT model. The model thus changes the CNTX variable more slowly at the higher levels compared to the lower levels. Of course, in addition to this fixed parameter *τ* which determines how much context is typically preserved in each level, the context updating is additionally influences by the surprise parameter, *α*, which can transiently increase the overwriting of prior context.

###### Information flow between levels of HAT is gated by surprise

We designed the feedforward information flow in HAT based on the notion that temporal integration is a distributed process, assuming that higher-level circuit perform a similar operation as lower-level circuits (i.e. linking input to prior context) but the higher levels may learn to associate chunks instead of single elements in the sequence. Our goal was that, for a multi-level compound like the word *airplane*, the first level of the model might learn to chunk the phonemes within *air* and *plane*, and the second level might learn to chunk *air* and *plane* to represent the larger word *airplane*. Thus, the input to the higher levels of the HAT model should be the compressed (chunked) representations from the lower levels. However, this process should also be modulated by surprise, as higher levels should only accept “successful” chunks from the layer below.

Therefore, the input to the higher levels of HAT is a linear mixture of HID and IN from the level below, modulated by the surprise *α*:

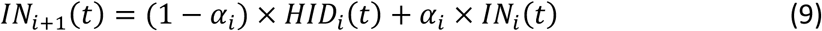

If the lower-level unit detects a large surprise (i.e. large *α*), more of the input from the lower level would be sent to the upper level as input. On the other hand, if the lower-level unit detects a small surprise (i.e. small *α*), more prior context or the “chunk” representation from the lower level would be preserved and sent to the upper level as input (Figure S2B).

### 2. Model Simulations and Predictions of Empirical Phenomena

#### 2.1 General Logic of Model Simulations

To demonstrate hierarchical context dependence in our models, we employ a strategy that is analogous to the original human experiments. We present the model with intact and scrambled versions of a time-varying stream of input. We then measure the context effects by comparing the model responses (internal representations) of the same input preceded by different contexts.

##### Training Procedure

We approximately simulated the experimental paradigm in Lerner et al. (2011) in order to examine whether each of our models exhibit a hierarchy of context dependence. We trained HAT, HAT-NR and TCM with a 30-element long “intact” sequence. The intact sequence was presented 600 times. Each element of the input was encoded as a one-hot vector of length 30. We also added uniformly distributed noise to each scalar value of each input sequence. The noise samples were independently drawn from a uniform distribution on [- 0.3, 0.3]. The purpose of the noise was to improve the model’s generalizability, and to approximate the fact that real-world sequence learning occurs in the presence of noise. To prevent the model from learning a spurious relationship between the end of the intact sequence and the beginning of the next presentation of the intact sequence, we added “random filler” sequences (length=5 symbols) between intact segments. Each of the random filler symbols was an independently generated random vector, with elements uniformly distributed in the range [-1, 1] (Figure S3A, depicted as an ‘x’ between intact segments).

##### Testing Procedure

After training, the weights in the model were frozen (no further weight change was allowed during test). We then compared the models’ representations of intact and scrambled sequences. The three scrambled sequences were designed to preserve the intact structure at three different scales: the long-scale (6 element subsequences were preserved), medium-scale (3 element subsequences were preserved) and fine-scale (2 element subsequences were preserved). Each testing ensemble (e.g. “medium scale scramble”) was composed of 10 “test sequences”. Each test sequence was a length-30 sequence which was a randomly scrambled version of the intact sequence. All test sequences within an ensemble were scrambled at the same scale, but with different permutations. Therefore, each test sequence exhibited preserved structure on the relevant scale. As during training, fillers were again inserted between each of the 10 sequences that composed a testing ensemble (Figure S3A, the ‘x’ between test sequences).

##### Quantifying Context Effects

We measured the similarity of the representations generated by different levels of the model as a function of the amount of shared context. To quantify similarity, we correlated the hidden representation that was generated when the models were processing the intact and the scrambled sequences (Figure S3A). Specifically, we correlated the hidden representations of the last elements of the subsequences (e.g. the “r” in “qr”) in each scrambled sequence with the hidden representations of the same elements (the “r” in “qr”) in the intact sequences (see red symbols in Figure S3A). In this way, we measured how the representation of the identical stimulus was altered as a function of the context change.

To assess whether the models really captured the temporal structure of the intact sequences due to sequence-specific learning (rather than due to an intrinsic ability to maintain prior context of any kind of sequence), we trained the models with random sequences generated by shuffling the intact structured sequences. We then tested the random-trained model with the same (non-random) testing sequences. We then quantified the context effects in the models in the same way as before, and could compare the context effects for the models trained with structured sequences against models trained with randomly shuffled sequences.

#### 2.2 Comparing the context dependence effects across model variants

We defined the “context dependence” (CD) effect as the difference in intact-scramble correlation across the long-scramble and short-scramble conditions: CD = corr(intact, LSS) – corr(intact, FSS).

To compare the CD in two models, we computed the distribution of CD values for each network and computed: (i) a t-test of the difference in means and (ii) a d-prime measurement of the separation of the distributions.

Training the HAT model on shuffled sequences produced a large and highly statistically significant decrease in sensitivity to temporal context. Comparing the original HAT model against the shuffle-trained variant, we obtained mean CD original = 0.25, mean CD shuffle- trained = 0.041, t(198) = 35, p < 0.001. In parallel, to characterize the effect size of this difference, we computed the Cohen’s d of the difference in distributions as d = 4.97.

#### 2.3 Model Performance in Generating Hierarchy of Context Dependence

As described in the main text, for a model to account for the hierarchy of context dependence it should capture two key phenomena:

(P1) lower processing stages of the model should be insensitive to context change (analogous to sensory cortical regions, Figure 1D, yellow bars);

(P2) increasingly higher processing stages of the model should be increasingly sensitive to temporal context further in the past (analogous to the higher stages of cortical processing, Figure 1D, blue bars).

##### Performance of Signal Gain Model

The signal gain model was able to account for both of the key empirical phenomena of hierarchical context dependence (P1 and P2, above). By manipulating the noise added to the internal representation which dependent on different levels of the processing stage and different levels of scrambled stimuli, the signal gain model generated the pattern of hierarchy of context dependence: the higher “stages” of the signal gain model generated lower correlation between intact and scrambled stimuli (Figure S3C, left). However, because the pattern was due to the noise added to the internal representation during testing, we observed a similar pattern when testing the model on temporal structures that it was not trained on (Figure S3C, right). In summary, signal gain model could account for the empirical data without preserving any temporal context.

##### Performance of TCM

Our passive integration model (hierarchical TCM) was able to account for both of the key empirical phenomena of hierarchical context dependence (P1 and P2, above). As expected, higher “stages” of the TCM model (lower *β* values) generated internal representations with greater temporal context dependence: this was manifest as a lower correlation between intact and scrambled stimuli (Figure S3D, left). TCM (and other linear integrator models) do exhibit an important failing, however: their context dependence was not selective for the temporal regularities that were presented during training. We observed a similar pattern of results when testing TCM on temporal structures that it was not trained on (Figure S3D, right). Thus, a “passive” form of temporal integration (exponential smoothing) as in the TCM model can account for much of the empirical data, although with some non-specificity for the kind of information that is being integrated.

##### Performance of HAT Model

Our active and gated integration hierarchical model (HAT) was able to account for both of the key empirical phenomena of hierarchical context dependence (P1 and P2, above). The lower levels of the model exhibited high correlation between representations of intact and scrambled stimuli, regardless of the scale of scrambling. This suggests that the lower level of the model is less sensitive to the change of the context. In contrast, the higher level of the model showed progressively more dependence on context: the correlation between the intact stimuli and the long-scale scrambled stimuli remained high, but the representation of the intact stimuli no longer resembles the medium scale or fine-scale representations (Figure S3E, left). Thus, the internal representations generated by the higher level of the model are sensitive to longer scales of temporal context.

The context dependence in the HAT model is not a generic property of its architecture, but depends on it having previously learned about the temporal regularities in the stimulus. Compared to the standard HAT model, a variant trained with shuffled sequences showed a context dependence effect that was less than 1/3 of the context dependence effect in the intact model (Figure S3E, right, see also Methods Section 2.2). There is still context dependence in this shuffle-trained variant, but this dependence likely arises mostly from the timescale parameters, *τ*, which are larger in higher order regions. Thus, the context dependence in this shuffle-trained model reflects the intrinsic rate of model updating, rather than the prior learning of temporal regularities.

##### Performance of HAT Variants

In order to test which ingredients of the HAT architecture are necessary for its functional properties, we tested a second variant of the HAT model which lacked a context gating procedure and also lacked a gradient of *τ* values. In this variant, HAT-NR (Methods Section 1.4), we observed lower intact-scramble correlation across all levels of the model, indicating that it has an inferior ability to represent the sequential structure of the stream of stimuli. Moreover, the lowest processing stage of this model already exhibited context dependence. In other words, the correlation between intact and long-scale scrambled stimuli was higher than the correlation between intact and fine-scale scrambled stimuli, even in the earliest stages of the HAT-NR model (Figure S3F, left). This is inconsistent with the empirical data (e.g. Figure 3A in Hasson et al., 2008; Figure 5 in Lerner et al., 2011). Thus, without the gating mechanism, the context dependence that we observed seemed to reflect the stage-by-stage hierarchical architecture of the model, rather than reflecting learning of the sequential structure in the data. Consistent with this interpretation, the HAT-NR model also exhibited similar results when it was trained with shuffled sequences (Figure S3C, right). This indicates that (as in the hierarchical TCM model) the temporal integration occurring in HAT-NR was arising from the dynamical timescales of each layer, and did not reflect the learning of temporal regularities on different scales.

#### 2.4 Modeling the construction and forgetting of temporal context

We set out to model the ISPC results using the HAT model and the TCM model. The training sequences and procedure were the same as for modeling of the Lerner et al. (2011) data. For testing, we only tested the models with the intact and the paragraph-level scrambled sequences (corresponding to the intact and scramble group of the empirical data). We added random noise sampled from a normal distribution ∼*N*(0, 3) to these two test models to generate the 100 simulated “subjects” for the intact and the scramble group, respectively. To better approximate the real neural data, we also included the effect of “hemodynamics” in our model, by convolving the timecourse of the hidden representations with a temporal smoothing function. The temporal smoothing function in the model consisted of two gamma functions to approximate the hemodynamic response, with peak value equal to 1 at t=1 and slowly decreasing to 0 at t=4.

Temporal context construction and forgetting were analyzed in the model in an analogous manner to how they were assessed in the empirical data. For clarity, we introduce notation that discriminates the cross-group similarity measure rSI for the context construction and context forgetting analyses. Specifically, we use rSI_ACCUMULATE_ = rSI_DE:CE_ for the rSI in the context construction analysis, and rSI_FORGET_ = rSI_CD:CE_ for the rSI in the context forgetting analysis. The context construction curve, rSI_DE:CE_, was estimated by computing ISPC on the internal representations of each model. Internal representations were measures as the same six-element segments were presented as input, preceded by different segments in the intact and scramble group. Similarly, for the forgetting curve, rSI_CD:CE_, the correlations were measured in the model simulation by performing ISPC on the hidden representations across two different “groups” of model runs. Each model run was treated in the same way as the neural response of a single participant. Thus, we measured the responses across two groups of model runs, where responses were correlated across different segments (e.g. segment D in Group 1 and segment E in Group 2) which were preceded by the same segment (e.g. segment C was the preceding segment in both Group 1 and Group 2).

### 3. Empirical Measurements: Constructing and Forgetting Temporal Context

#### 3.1 Subjects

Forty-four subjects (all native English speakers) were recruited from the Princeton community (20 male, 24 female, ages 18-29) and nine subjects (all native English speakers) were recruited from the Johns Hopkins community (5 male, 4 female, ages 19-41). Conditions in which the head motion were >1 mm or where the signal was corrupted were discarded from the analysis. Overall, 31 subjects were participated in the intact condition, and 31 subjects were participated in the scramble condition. All subjects had normal hearing and provided written informed consent.

#### 3.2 Stimuli and Experimental Design

Stimuli for the experiment were generated from the 9 min real-life story (“It’s not the fall that gets you,” told by Andy Christie) recorded at a live storytelling performance (“The Moth” storytelling event, New York City). Subjects listened to the whole story from beginning to end (intact forward story), as well as to the scrambled stimuli, which were created by randomly shuffling segments of the intact story. More specifically, the story was segmented manually by identifying the end points of the 25 segments, and was randomly scrambled by these segments (21.9 ± 4.29s). The two conditions both started with a 25 seconds intro music, which were discarded from all analyses. For the subjects who listened to both conditions, the scrambled condition was played first before the intact condition, to reduce the influence of prior knowledge of the story.

#### 3.3 Preprocessing of Neuroimaging Data

Imaging data were acquired on a 3T full-body scanner (Siemens Skyra for data from Princeton; Phillips Elition for data from Johns Hopkins University) with a 20-channel head coil using a T2*- weighted echo planar imaging (EPI) pulse sequence (TR 1500 ms, TE 28 ms, flip angle 64, whole-brain coverage 27 slices of 4 mm thickness, in-plane resolution 3 by 3 mm, FOV 192 by 192 mm). Preprocessing was performed in FSL, including slice time correction, motion correction, linear detrending, high-pass filtering (140 s cutoff), and coregistration and affine transformation of the functional volumes to a template brain (MNI). Functional images were resampled to 3 mm isotropic voxels for all analyses.

#### 3.4 Inter-subject pattern correlation (ISPC) analysis

The ISPC analysis quantifies the similarity of spatial patterns of neural responses at a moment in time. We quantify the similarity by correlating the pattern of voxel activation at each time point (Figure 2A). Similar to the inter-subject correlation (ISC) analysis which provides a measure of the temporal reliability of the responses to complex stimuli (Hasson et al. 2009), the ISPC analysis provides a measure of the spatial reliability of the response to the stimuli at each time point. The ISPC method differs from conventional fMRI data analysis methods in that it circumvents the need to specify a model for the neuronal processes in any given brain region during story listening. Instead, the ISPC method uses one subject’s neural responses to a stimulus as a model to predict the neural responses within other subjects.

Using ISPC, quantified the changes in the neural responses over time within each segment of the auditory stimulus. We computed similarity within the group of subjects listening to the intact story (the intact condition, rII), similarity within the group listening to the scrambled story (the scramble condition, rSS), and similarity across the intact and scrambled groups (rSI) (Figure 2B). The rII and rSS analyses provide an indication of how reliably a given region is responding to the stimulus (Intact or Scrambled) at a particular moment. Conversely, the rSI analysis across the two groups indicates the similarity across two groups, which may be experiencing the same input (but different contexts) or experiencing different input (but with the same prior context). For example, the main analysis (Figure 2C) examines the similarity across intact and scrambled groups when subjects process the same segments preceded by different contexts: we correlated responses to segment E, which was preceded by segment D in the intact group but preceded by segment C in the scrambled group). That is, when the context is disrupted in the scramble group, we measure how subjects re-construct the temporal context in order to align with the intact group.

#### 3.5 Procedure for calculating similarity within groups (rII and rSS) and between groups (rSI)

To calculate rII, we segmented the neural response according to the segments used to make the scrambled stimuli. For each segmentation, we analyzed the neural response of the first 16 seconds. We performed ISPC by correlating the neural response pattern of one subject in the intact group to the average neural responses of the remaining subjects in the intact group for each time point. This calculation of spatial patterns was performed separately for each timepoint in each segment. We generated an ISPC time course within each long segment for each subject. The rII was calculated by averaging the ISPC time course across all segments and subjects. The rSS was calculated using the same method, within the scramble group. To calculate rSI, for each long segment, we performed ISPC by correlating the neural response pattern of one subject in the scramble group to the average neural response of all subjects in the intact group. The rSI time-course was calculated by averaging the ISPC time-course across all segments and all subjects.

#### 3.6 Regional pattern analysis: construction of temporal context

To discriminate the three possible models (See Result section), we first analyzed ISPC in the primary auditory cortex (A1) and the right temporal parietal junction (rTPJ). To more precisely partition the neural responses into distinct segments, we up-sampled the neural response to 50Hz. Furthermore, the neural response time courses of all ROIs were aligned across all subjects based on the signal of A1, and then aligned with the audio waveform of the stimuli (r=0.31). After the alignment of the signal, we divided the story into segments and performed the ISPC analysis as described in the previous paragraph.

To further examine whether there is a hierarchy of context construction along the cortical hierarchy, we conducted the ISPC analysis for 400 ROIs across the whole brain, based on the parcellation of the cerebral cortex provided by Schaefer et al., (2018). We only analyzed ROIs that showed reliable responses across subjects listening to the scrambled stimuli. For selecting ROIs, we chose an arbitrary threshold of mean(rSS) > 0.06 over all time points, which produces a set of ROIs which corresponds to prior ISC maps (See Methods Section 3.7, Figure S5). To quantify the rise time of different regions, we fit the rSI_DE:CE_ curves with the logistic function

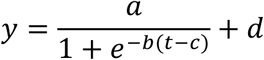

by using least-square regression to minimize error. Here, *y* is the dependent variable which is the rSI value and *t* is the time in seconds since the segment onset. The fitted parameter *a* is the curve’s maximum value, *b* is the steepness of the curve, *c* is the time when the logistic curve reaches its half maximum value, and *d* is an offset term to adjust the initial value of the curve. Among the 83 ROIs which exhibited reliable responses to the scrambled stimuli, 4 ROIs were excluded because the parameter of the logistic function could not be recovered with confidence after fitting the rSI curves (Figure S6) and 79 ROIs were proceeded to final rise time analysis (Figure S7). The rise time for the intact-scramble alignment was defined as the time when the logistic curve reaches its half maximum value (i.e. parameter *c*). The rise time of the individual ROIs were mapped from MNI space to a cortical space, and visualized on a cortical surface map using Workbench Viewer (https://www.humanconnectome.org/software/connectome-workbench).

#### 3.7 Validating the reliability of neural responses when subjects are listening to the scrambled stimuli

To determine the ROIs that showed reliable responses when people were processing the naturalistic narratives, we first chose an arbitrary threshold for rSS = 0.06 which produced a set of ROIs which corresponds to prior ISC maps showed in Lerner et al. (2011). We further validated the threshold by conducting a permutation test of the rII in primary auditory cortex, in which we compared the true rII with the “shuffled rII” calculated after shuffling the order of the segments: We first generated 10000 shuffled orders of the segments. For each of these shuffled orders, we reordered the neural responses of one subject according to the shuffled order, and calculate the rII between the shuffled neural response of this one subjects with the mean neural responses of the rest of the subjects in the intact group (For the other subjects, the order was preserved). We repeated this procedure to all the 31 subjects in the intact group using the same 10000 shuffled orders, and calculated the average rII for each shuffled order, generating a null distribution of the rII. We found that 0.06 is significantly higher than the null distribution (p<0.0001, Figure S5), confirming that this is a valid threshold for determining ROIs showing reliable responses when people are processing naturalistic stimuli. Furthermore, the raw curves of rII and rSS showed that the ROIs showed reliable responses from the beginning to the end of the segments (Figure 2F). This indicates that the ROIs determined by rSS=0.06 showed meaningful responses when people were processing the intact and the scrambled stimuli.

#### 3.8 Regional pattern analysis: forgetting of temporal context

To test the predictions of the HAT and TCM models (See *Modeling the temporal context construction and forgetting*), we performed another ISPC analysis to examine the temporal context forgetting. To do so, we examined the similarity of neural responses over time across two groups of subjects processing the different segments preceded by the same context. For example, in the rSI_FORGET_ (= rSI_CD:CE_) analysis, we correlated the responses between segment D in the intact group and segment E in the scramble group, when both were preceded by segment C (Figure 4B). The procedure for calculating procedure of rSI_FORGET_ (= rSI_CD:CE_) was directly analogous to the calculation of rSI_CONSTRUCT_ (as illustrated in Figure 2) except that we paired non-matching segments with matching contexts (CD:CE), rather than pairing matching segments with non-matching contexts (DE:CE).

The HAT model exhibited variable rSI_FORGET_ curves across simulation runs, because network weight initialization and the randomization order for scrambling affect the hidden representations that are learned, and how these drive context gating. Therefore, we plot a representative rSI_FORGET_ simulation in Figure 4G. Despite this variability, the HAT model’s behavior was consistently different from the TCM model, which predicts that regions with slower changes in rSI_CONSTRUCT_ will also exhibit slower changes in rSI_FORGET_.

