## Supplemental Figures and Text for "Constructing and Forgetting Temporal Context in the Human Cerebral Cortex"

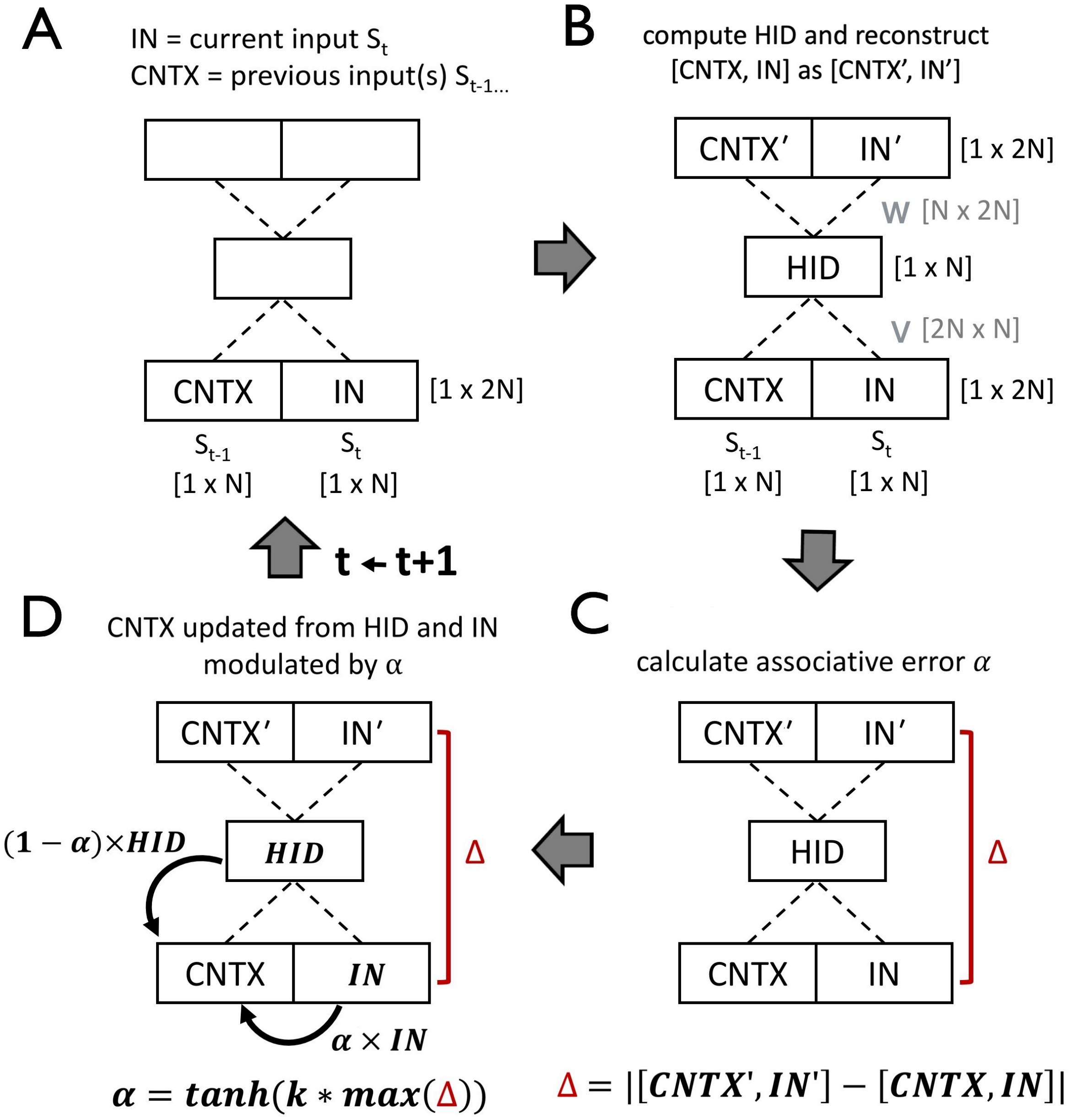


**Figure S1. Architecture and updating of a local autoencoder-in-time (AT) unit. (A)** At time t, the input layer of the AT unit is a 1-by-2N vector which contains both the present information $S_t$ in the IN bank and the past information $S_{t-1}$ in the CNTX bank. (B) The concatenated vector [CNTX, IN] is multiplied by weight matrix V to form a low-dimensional HID representation (a 1-by-N vector). This HID vector is then left-multiplied by a weight matrix W to generate an output layer [CNTX′, IN′] which is the reconstruction of input [CNTX, IN]. (C) The reconstruction error, Δ, or “surprise”, is calculated as the absolute value of [CNTX′, IN′]- [CNTX, IN]. (D) The gating parameter, α, is then calculated as tanh(k*max(Δ)). Here, the parameter k scales how much the contribution of IN to CNTX is increased by surprise. The CNTX vector is updated as a linear mixture of the IN vector and HID vector, with the linear proportions modulated by α and a level-specific time constant τ. After CNTX is updated, the cycle is complete, and the unit is ready to receive input at time (t+1). IN = input unit, CNTX = context unit, HID = hidden state unit.

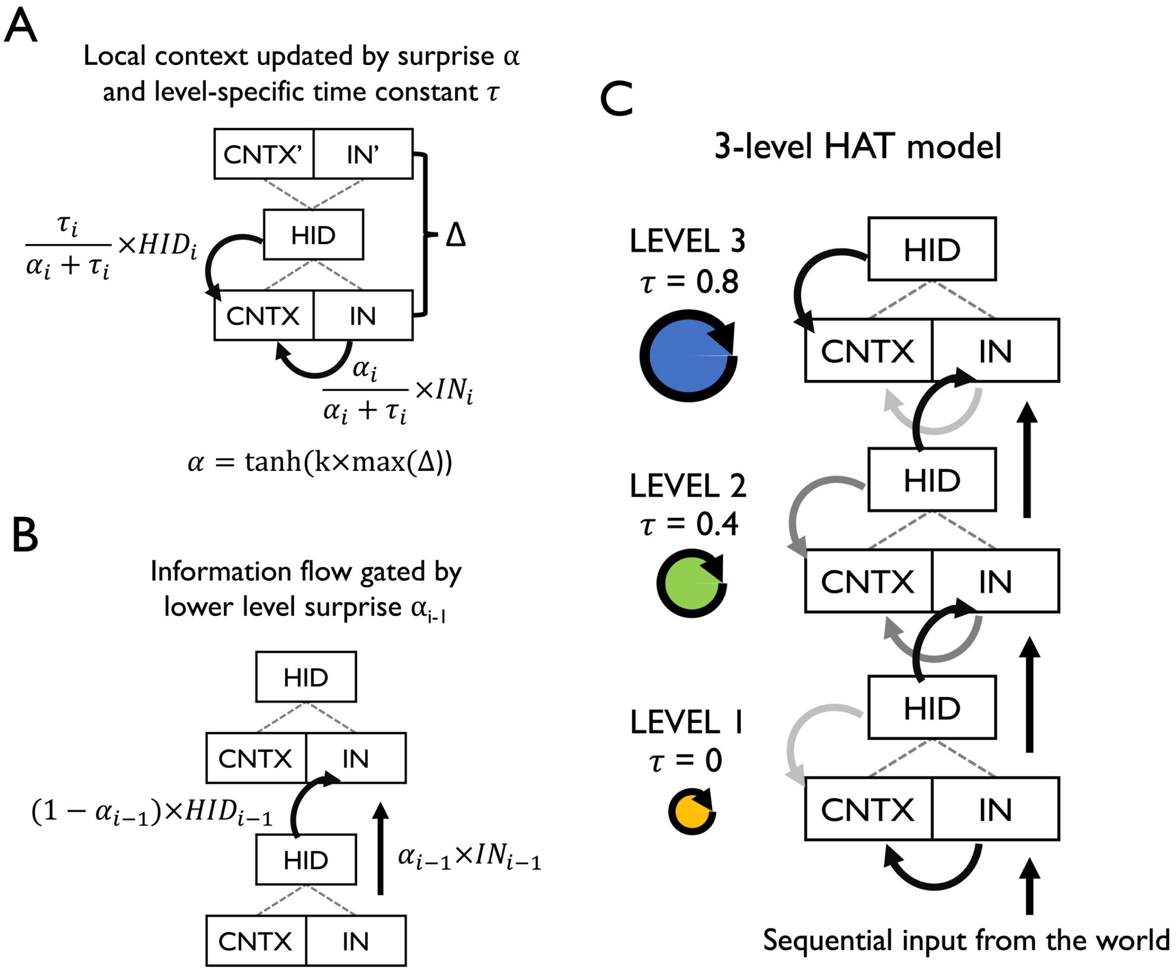


**Figure S2**. **Architecture and information flow of the hierarchical autoencoder in time (HAT) model.** (A) An autoencoder in time (AT) unit, in which the local context CNTX is updated with hidden representation HID and current input IN, modulated by specified time constant τ and "surprise" α. α is computed via auto-associative error Δ and a scaling parameter k. If τ is larger than α, the model unit tends to preserve more context from HID. Otherwise, the model unit tends to overwrite the context with the current input IN. (B) Information to level i is gated by surprise α from level (i-1). If $\alpha_{i-1}$ is large, more information from $IN_{i-1}$ is sent to level i. If $\alpha_{i-1}$ is small, then more information from $HID_{i-1}$ (i.e. more history-dependent information) is sent to level i. (C) A three-level HAT model. Each level is an AT unit, which approximates a single stage of cortical processing. Each level was assigned a time constant, τ, which determines the amount of local context that is preserved (on average) at that level of processing (see panel A). Higher levels of the model were assigned larger τ values than lower levels of the model. The input to Level 1 is the external sequential stimuli. The input to the higher levels is a linear mixture of IN and HID from the level below, modulated by α (see panel B). IN = input unit, CNTX = context unit, HID = hidden state unit, HAT = hierarchical autoencoders in time.

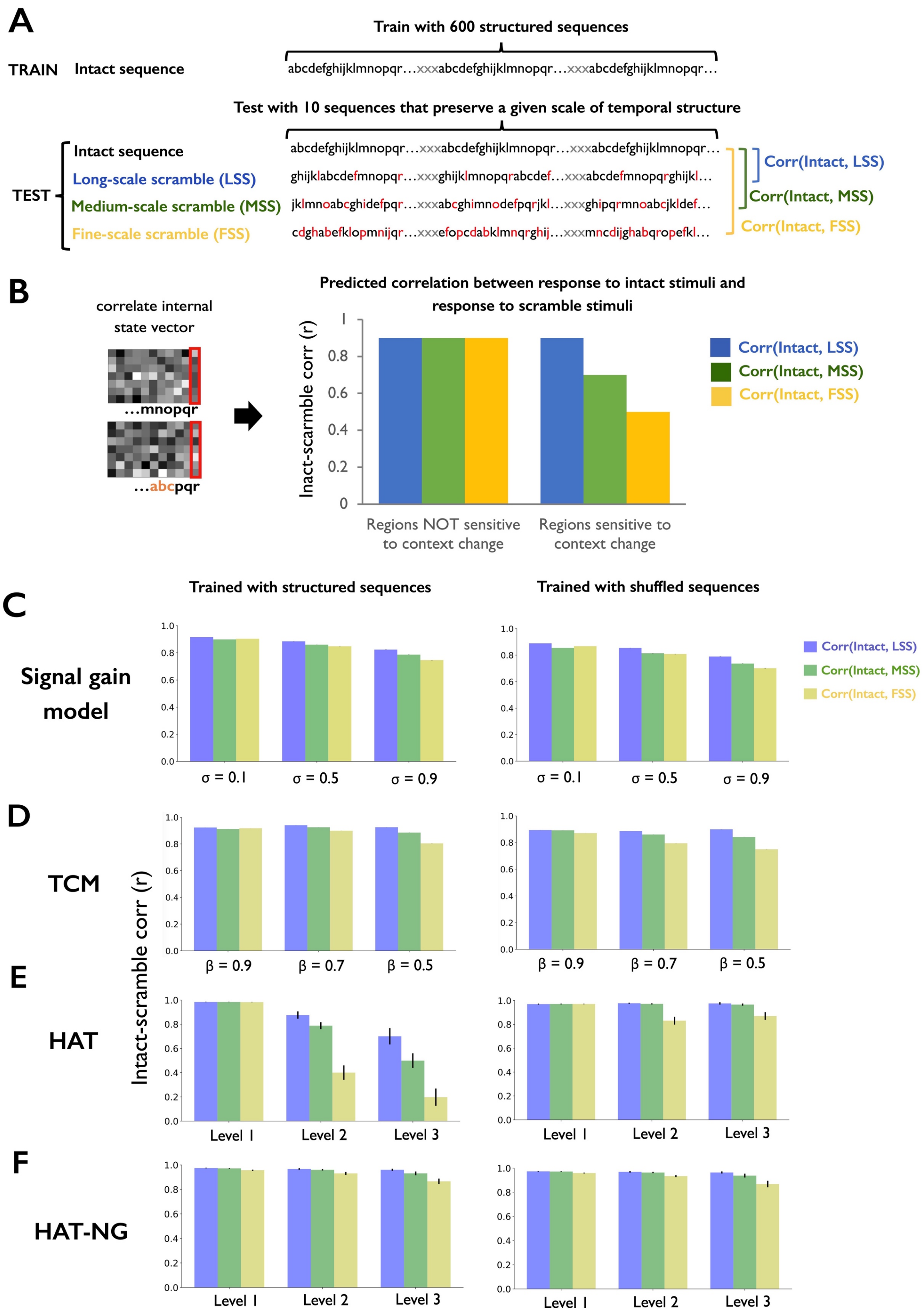
A
Train with 600 structured sequences
TRAIN Intact sequence
abcdefghijklmnopqr…xxxabcdefghijklmnopqr…xxxabcdefghijklmnopqr…
Test with 10 sequences that preserve a given scale of temporal structure
TEST
Intact sequence
Long-scale scramble (LSS)
Medium-scale scramble (MSS)
Fine-scale scramble (FSS)
abcdefghijklmnopqr…xxxabcdefghijklmnopqr…xxxabcdefghijklmnopqr…
ghijklabcdefmnopqr…xxxghijklmnopqrabcdef…xxxabcdefmnopqrghijkl…
jklmnoabcghidefpqr…xxxabcghimnodefpqrjkl…xxxghipqrmnoabcjkldef…
cdghabefklopmnijqr…xxxefopcdabklmnqrghij…xxxmncdijghabqropefkl…
Corr(Intact, LSS)
Corr(Intact, MSS)
Corr(Intact, FSS)
B
correlate internal state vector
…mnopqr
…abcpqr
Predicted correlation between response to intact stimuli and response to scramble stimuli
Inact-scramble corr (r)
Regions NOT sensitive to context change
Regions sensitive to context change
Corr(Intact, LSS)
Corr(Intact, MSS)
Corr(Intact, FSS)
C
Trained with structured sequences
Trained with shuffled sequences
Signal gain model
σ = 0.1
σ = 0.5
σ = 0.9
Corr(Intact, LSS)
Corr(Intact, MSS)
Corr(Intact, FSS)
D
TCM
β = 0.9
β = 0.7
β = 0.5
E
HAT
Intact-scramble corr (r)
Level 1
Level 2
Level 3
F
HAT-NG
Level 1
Level 2
Level 3

**Figure S3. The signal gain model, passive integration model (TCM) and active integration model (HAT) can account for prior data on hierarchical context dependence. (A)** Example of training sequences (intact sequences) and testing sequences (long scale, medium scale and fine scale scrambled sequences). Context dependence was measured by correlating the hidden representation between the intact and different levels of scrambled sequences. The target element (i.e. the last element of each sub-sequence) for correlation is marked with red. **(B)** The predicted correlation of hidden representations in regions that are more / less sensitive to temporal context. **(C)** The signal gain model showed more sensitivity to different levels of the context change when adding more noise on higher level stages of the model. This effect was not specific to testing with any particular sequence structure. **(D)** TCM showed more sensitivity to different levels of the context change when the $\beta$ parameter was decreased (i.e. when the model preserved more temporal context, analogous to the higher-level circuit). However, this context-dependence effect in TCM was not specific to sequences that were seen during training – it was also observed when training and testing employed completely different sequences. **(E)** HAT trained with structured sequences showed a hierarchy of context dependency across different levels of the model. Importantly, this context dependence effect in HAT was much stronger when the model was trained and tested on the same sequences. In other words, the context dependence in HAT depends on the model's learning of temporal structure. **(F)** HAT without a gating mechanism (HAT-NG) showed a similar pattern to the TCM and signal gain results: the higher levels of the model showed more context dependence, but the pattern generated was not specific to the structure of the training sequences. LSS = long scale scramble, MSS = medium scale scramble, FSS = fine scale scramble. TCM = temporal context model. HAT = hierarchical autoencoders in time. HAT-NG = HAT with no gating mechanism.

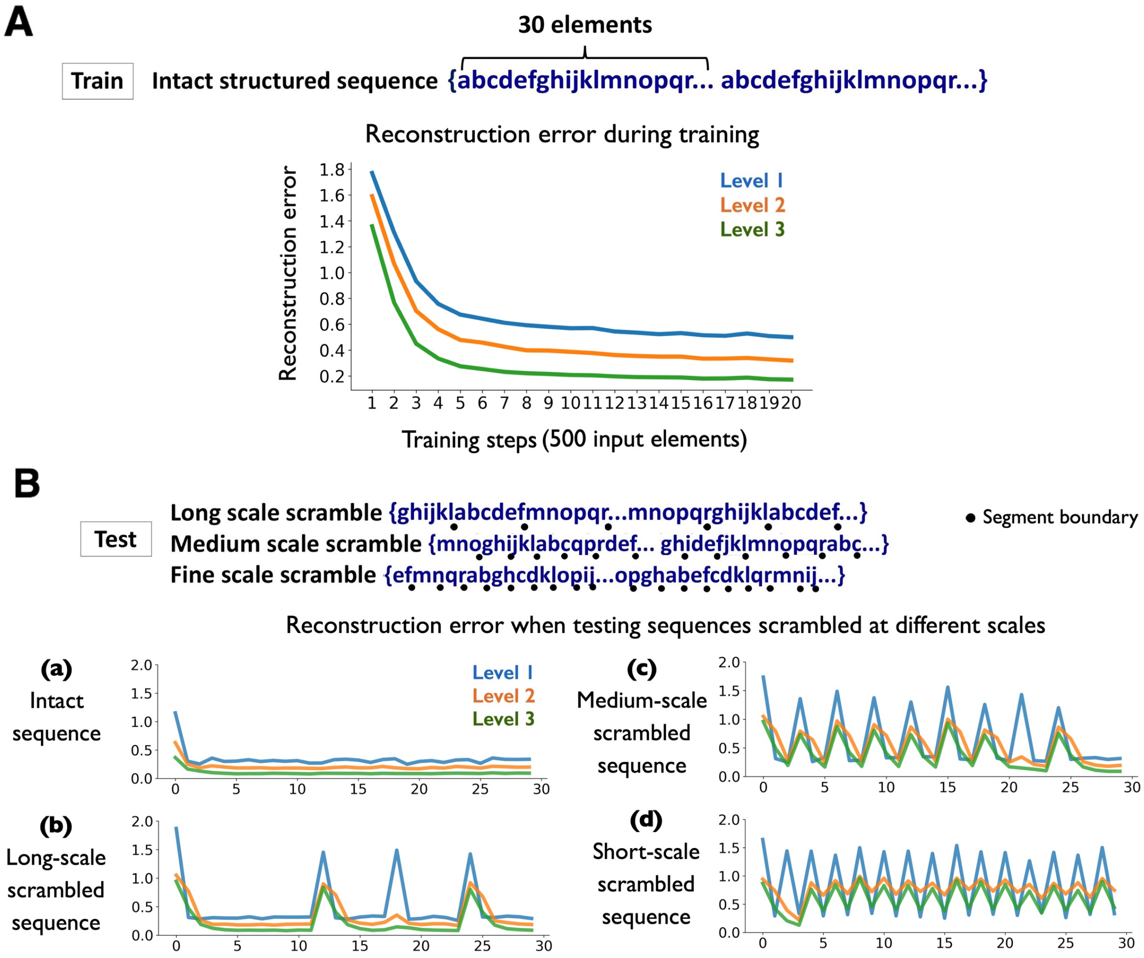


**Figure S4**. **Training and testing error generated by the HAT model.** (A) During training with the structured sequences, all levels of HAT showed decreasing reconstruction error Δ as training duration increased. (B) Reconstruction error when testing with different levels of scrambled sequences: (a) intact sequence (b) long-scale scrambled sequences (c) medium-scale scrambled sequences (d) short-scale scrambled sequences. HAT generated higher error when detecting sequence boundaries based on different scales of scrambling.

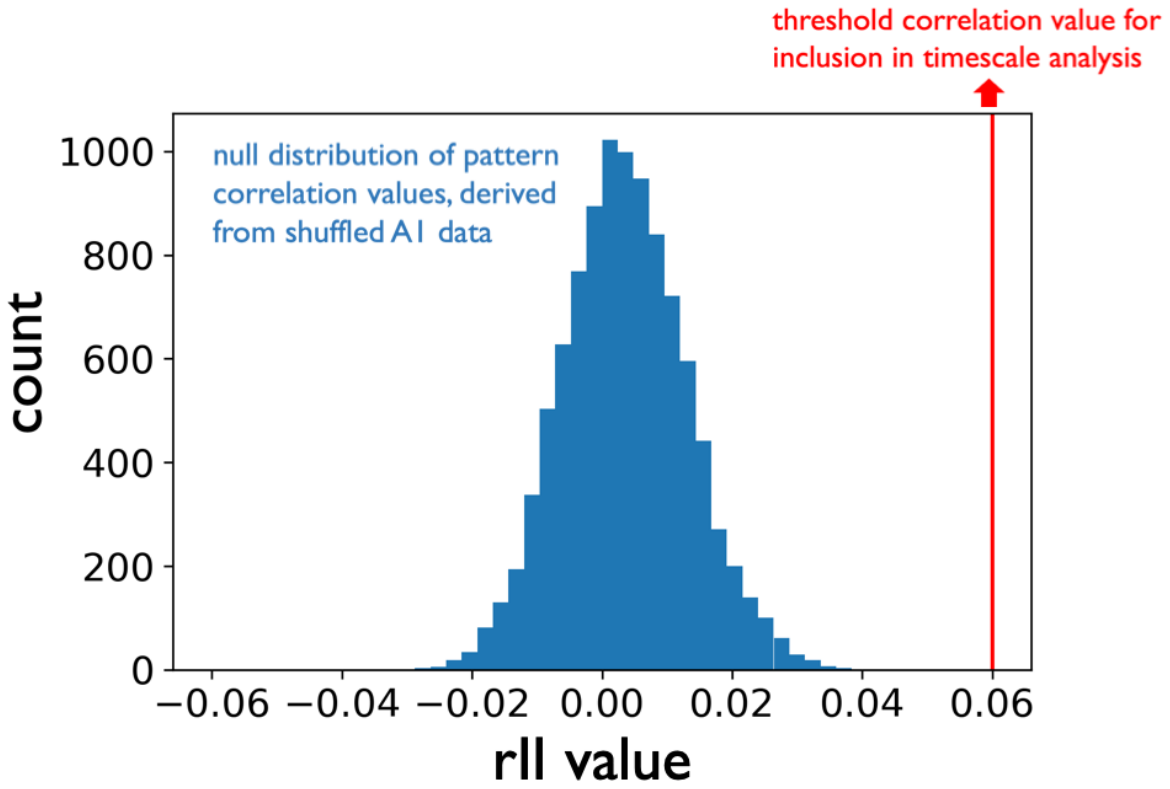


**Figure S5. Empirical rII value for auditory cortex is plotted in reference to a null distribution of rII values.** The within-group ISPC (rII) was computed within an auditory cortex "A1+" parcel, which was functionally defined in a separate naturalistic narrative dataset (Simony et al., 2016). The surrogate distribution of rII values was computed by computing ISPC against non-matching sentences (shuffling the sentence order, see Supplemental Methods Section 4). In order to visualize the most meaningful timescale parameters in regions that responding reliably (in Figures 2 and 3), we chose a threshold of rII=0.06. This threshold was not chosen in order to correspond to an arbitrary statistical threshold, but nonetheless it is clear that rII= 0.06 lies far outside the null distribution of rII values. Thus, we used 0.06 as a conservative threshold for ROIs that showed reliable stimulus-locked response. The ROIs included in Figures 2, 3 and 4 (all possessing rUU > 0.06) reflect a reliable stimulus-locked response to the scrambled stimulus. A1 = primary auditory cortex, rII = intact-intact inter-subject pattern correlation.

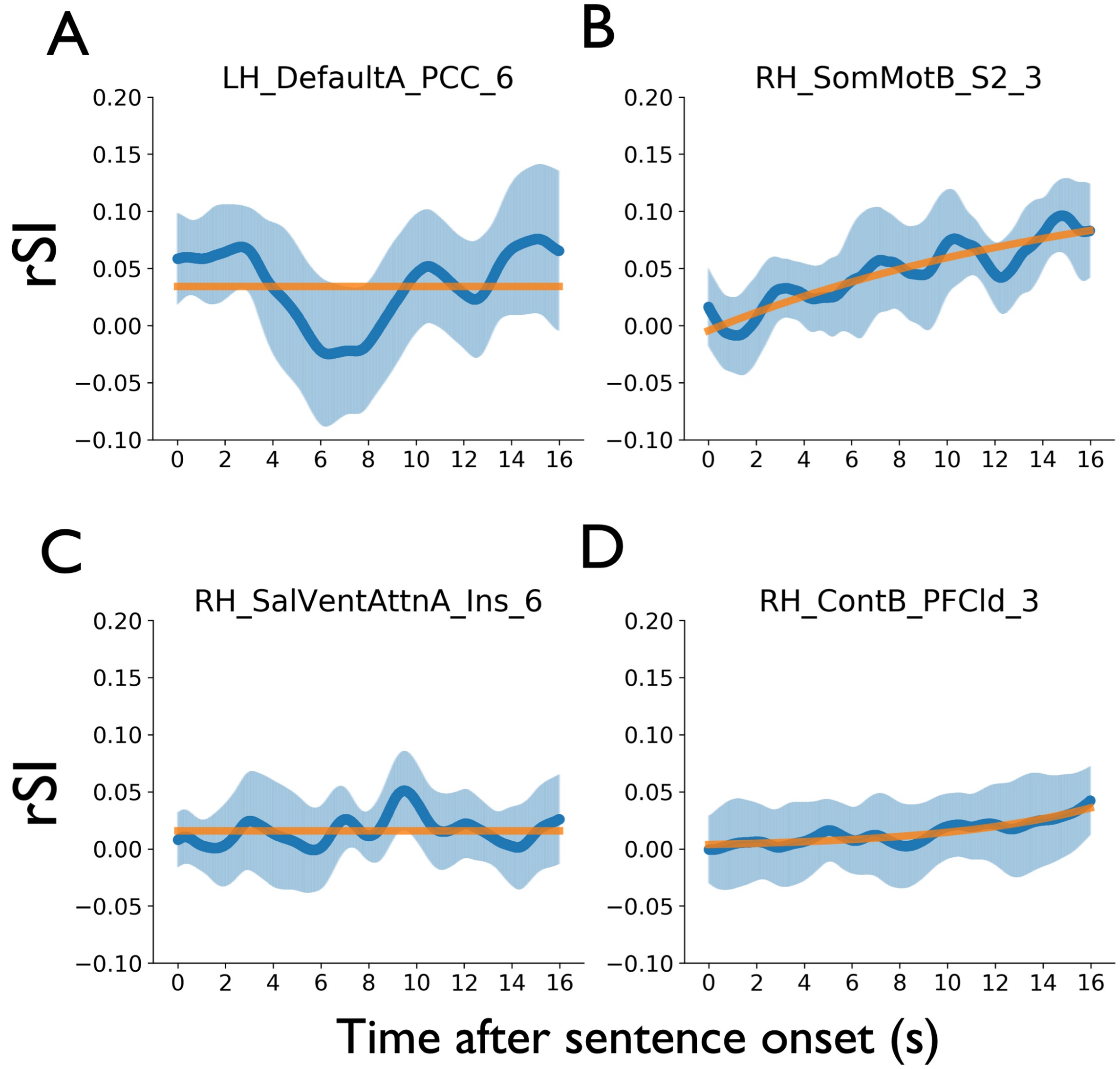


**Figure S6. A set of 4 anatomical regions of interest (ROIs) in which the parameters of the logistic function could not be confidently recovered after fitting the rSI curves.** We visually identified parcels in which the rSI curve did not appear to follow a logistic curve. The parcels are individually labeled with their names from the Schaefer parcellation (Schaefer et al., 2018). These parcels occur, (A) near left posterior cingulate cortex; (B) right somatomotor cortex; (C) right insula and (D) the right prefrontal cortex. rSI = scramble-intact inter-subject pattern correlation.

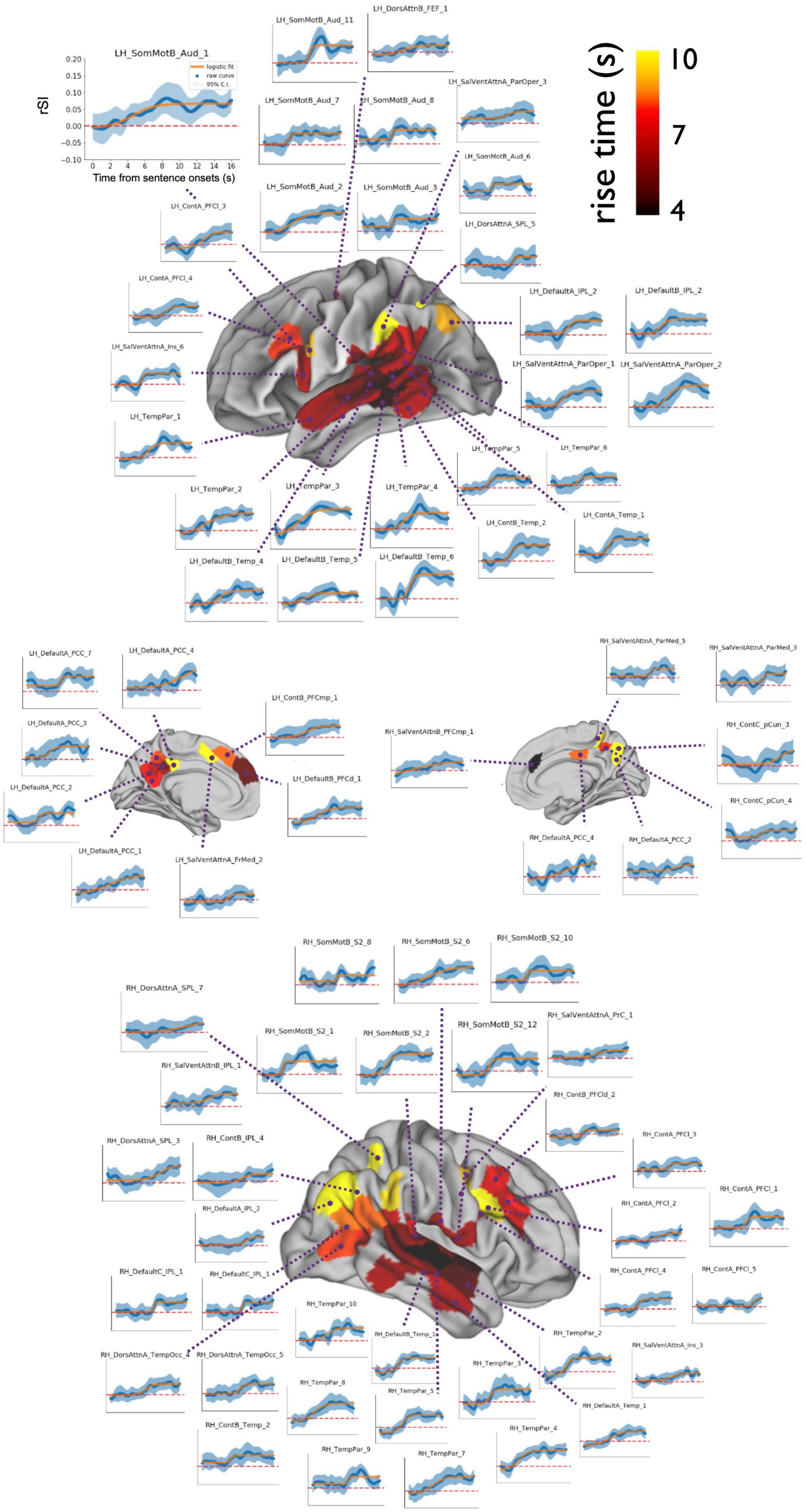
LH_SomMotB_Aud_1
logistic fit
raw curve
95% C.I.
rSI
Time from sentence onsets (s)
rise time (s)
10
7
4
LH_SomMotB_Aud_11
LH_DorsAttnB_FEF_1
LH_SomMotB_Aud_7
LH_SomMotB_Aud_8
LH_SalVentAttnA_ParOper_3
LH_SomMotB_Aud_6
LH_SomMotB_Aud_2
LH_SomMotB_Aud_3
LH_DorsAttnB_SPL_5
LH_ContA_PFCl_3
LH_ContA_PFCl_4
LH_DefaultA_IPL_2
LH_DefaultB_IPL_2
LH_SalVentAttnA_Ins_6
LH_SalVentAttnA_ParOper_1
LH_SalVentAttnA_ParOper_2
LH_TempPar_1
LH_TempPar_5
LH_TempPar_6
LH_TempPar_2
LH_TempPar_3
LH_TempPar_4
LH_ContA_Temp_1
LH_ContB_Temp_2
LH_DefaultB_Temp_4
LH_DefaultB_Temp_5
LH_DefaultB_Temp_6
LH_DefaultA_PCC_7
LH_DefaultA_PCC_4
LH_ContB_PFCmp_1
LH_DefaultA_PCC_3
LH_DefaultA_PCC_2
LH_DefaultB_PFCd_1
LH_DefaultA_PCC_1
LH_SalVentAttnA_FrMed_2
RH_SalVentAttnA_ParMed_5
RH_SalVentAttnA_ParMed_3
RH_SalVentAttnB_PFCmp_1
RH_ContC_pCun_3
RH_ContC_pCun_4
RH_DefaultA_PCC_4
RH_DefaultA_PCC_2
RH_SomMotB_S2_8
RH_SomMotB_S2_6
RH_SomMotB_S2_10
RH_DorsAttnA_SPL_7
RH_SomMotB_S2_1
RH_SomMotB_S2_2
RH_SomMotB_S2_12
RH_SalVentAttnA_PrC_1
RH_SalVentAttnB_IPL_1
RH_ContB_PFCld_2
RH_ContA_PFCl_3
RH_DorsAttnA_SPL_3
RH_ContB_IPL_4
RH_DefaultA_IPL_2
RH_ContA_PFCl_2
RH_ContA_PFCl_1
RH_DefaultC_IPL_1
RH_DefaultC_IPL_1
RH_ContA_PFCl_4
RH_ContA_PFCl_5
RH_TempPar_10
RH_DefaultB_Temp_1
RH_TempPar_2
RH_SalVentAttnA_Ins_3
RH_DorsAttnA_TempOcc_4
RH_DorsAttnA_TempOcc_5
RH_TempPar_3
RH_TempPar_8
RH_TempPar_5
RH_DefaultA_Temp_1
RH_ContB_Temp_2
RH_TempPar_4
RH_TempPar_9
RH_TempPar_7

**Figure S7. The temporal profiles of context construction mapped for each ROI individually.** The raw $rSI_{DE:CE}$ curves (blue curves) are overlaid with their corresponding logistic fits (orange lines) for each ROI. The shaded blue area indicates a parametric 95% confidence interval on each rSI measurement at each time point. rSI = scramble-intact inter-subject pattern correlation.

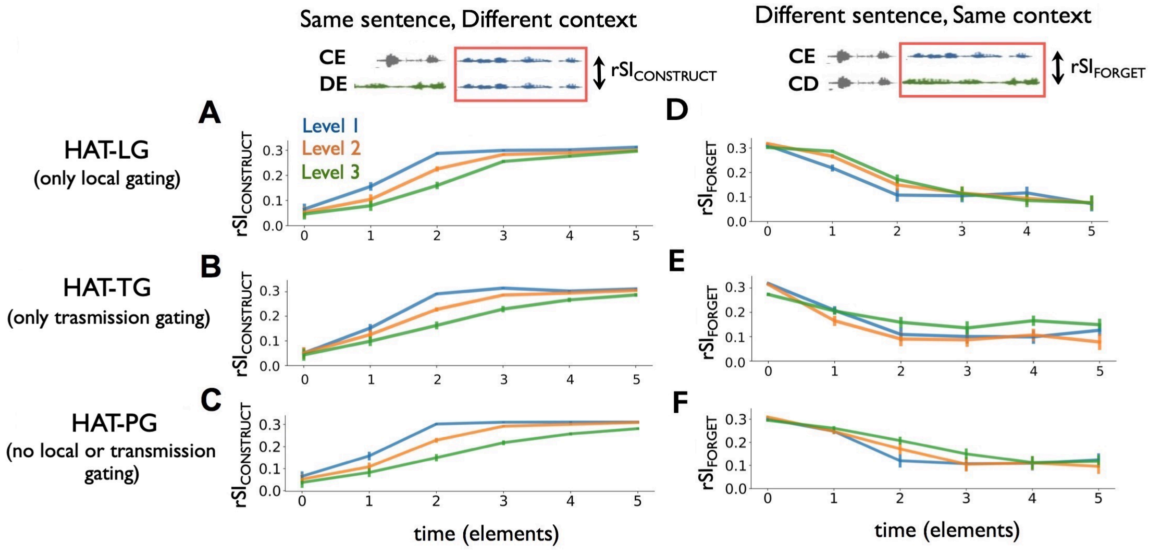


**Figure S8. Predictions of context construction ($rSI_{CONSTRUCT}$) and context forgetting ($rSI_{FORGET}$) for variants of the HAT model with limited gating mechanisms.** Each of these HAT variants has a limited gating mechanism (Supplemental Methods). For the context construction analysis: (A) Predictions of $rSI_{CONSTRUCT}$ generated by HAT with only local gating. (B) Predictions of $rSI_{CONSTRUCT}$ generated by HAT with only transmission gating. (C) Predictions of $rSI_{CONTRUCT}$ generated by HAT with no local gating and no transmission gating. For the context forgetting analysis: (D) Predictions of $rSI_{FORGET}$ generated by a model with only local gating. (E) Predictions of $rSI_{FORGET}$ generated by a model with only transmission gating. (F) Predictions of the $rSI_{FORGET}$ curve generated by model with no local gating and no transmission gating. HAT-LG = HAT with only local gating, HAT-TG = HAT with only transmission gating, HAT-PG = HAT with partial gating, rSI = scramble-intact inter-subject pattern correlation.

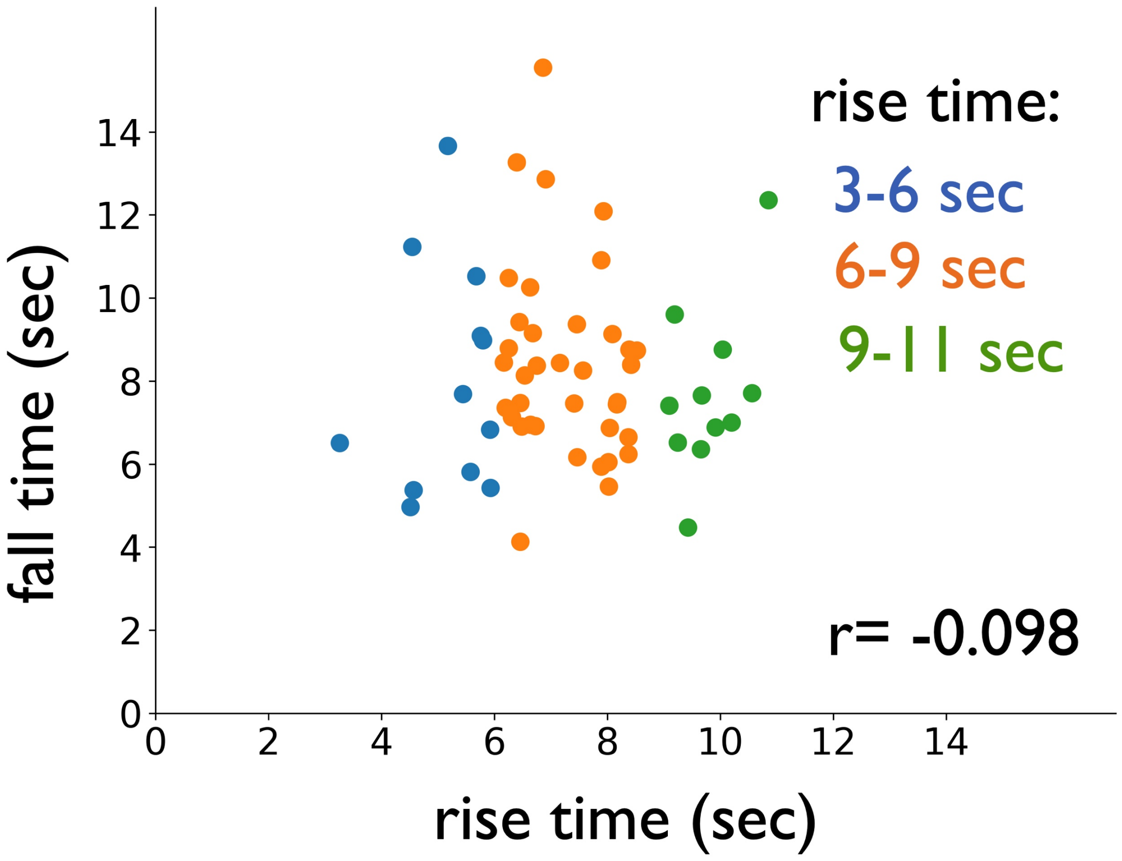


**Figure S9. The rise time (time for integrating prior information) does not match the forget time (time for forgetting prior information).** For each individual ROI, the rise time for context construction is plotted against the fall time for context forgetting in that ROI. There is no significant correlation (r=-0.1, p=0.5)

**Variants of HAT model with limited reset mechanisms**

To investigate which elements of the HAT model were necessary for generating empirical phenomenon, we generated a set of HAT models with no or only partial gating of temporal context. Specifically, we turned off the surprise-modulated context gating mechanism, either locally (i.e. the within-level context gating) or globally (i.e. the between-level transmission gating), or we turned off all gating effects.

**HAT-NG**

HAT-No Gating or HAT-NG, is a HAT model without any gating mechanisms. Locally, we only update the context with the hidden representation, by setting $\alpha$ in equation (6, main text) equal to 0:

$$CNTX_i(t+1) = HID_i(t) \tag{1}$$

Therefore, the $\tau$ gradient has no effect in the HAT-NR model.

Globally, we turned off the surprise-modulated transmission gating by setting $\alpha$ in equation (9, main text) equal to 0, so that the input of the upper level is purely the HID from the lower level. I.e.

$$IN_{i+1}(t) = HID_i(t) \tag{2}$$

**HAT-PG**

HAT-Partial Gating or HAT-PG, is a HAT model with a partial local gating mechanism. There is no surprise or $\alpha$ modulated gating mechanism; instead, the local context is reset by a fixed amount of input based on the level-specific $\tau$.

$$CNTX_i(t+1) = \tau_i \times HID_i(t) + (1-\tau_i) \times IN_i(t) \tag{3}$$

HAT-PG has no global gating. The input of the upper level is purely the HID from the lower level, as described in equation (2).

**HAT-LG**

HAT-Local Gating or HAT-LG, is a HAT model with no global gating. i.e. the input to the higher level of the model is simply a copy of the HID from the lower level, regardless of the $\alpha$ parameter (as in equation 2). However, the within-level CNTX update is still gated by surprise (equation 8, main text).

**HAT-TG**

HAT-Transmission Gating or HAT-TG, is a HAT model with no local gating, i.e. there is no $\alpha$ modulated gating mechanism. The local context is reset by a fixed amount of input based on the level-specific $\tau$ (equation 3). However, the between-level transmission is still gated by surprise as in equation (9, main text).
